# Tumor-specific CD4 T cells instruct monocyte differentiation in pancreatic ductal adenocarcinoma

**DOI:** 10.1101/2022.11.09.515858

**Authors:** Michael T. Patterson, Adam L. Burrack, Yingzheng Xu, Zoe C. Schmiechen, Patricia R. Schrank, Ainsley E. Kennedy, Maria M. Firulyova, Ebony A. Miller, Eduardo Cruz-Hinojoza, Konstantin Zaitsev, Jesse W. Williams, Ingunn M. Stromnes

## Abstract

Pancreatic ductal adenocarcinoma (PDA) is a lethal malignancy resistant to immunotherapy. The pancreatic tumor microenvironment is shaped and maintained by myeloid cells that outnumber tumor cells. Here, using monocyte fate-mapping PDA mouse models and human tumor tissues, we identify monocytes give rise to most heterogeneous macrophage subpopulations in PDA. We show that monocyte differentiation is governed by the local presence of CD4, but not CD8, T cells. We demonstrate that tumor specific CD4 T cells induce monocyte differentiation into antitumor MHCII^hi^ proinflammatory macrophages dependent on non-redundant IFNγ and CD40 signaling pathways that suppress tumor growth. Pancreatic tissue-resident macrophages exhibit an immunosuppressive pro-tumor state that is refractory to the modulatory effects of antitumor CD4 T cells. Intratumoral monocytes adopt a pro-tumor fate indistinguishable from tissue-resident macrophages following CD4 T cell depletion. Thus, tumor-specific CD4 T cell governance of monocyte fate promotes immune-mediated control of solid tumors.

**Highlights:**

▪ Circulating monocytes are progenitors to most heterogeneous macrophage subsets in PDA
▪ Monocyte-derived macrophage acquisition of an MHCII^hi^ phenotype is dependent on tumor-specific CD4 T cells
▪ In the absence of CD4 T cells, monocyte-derived macrophages acquire tissue resident macrophage traits and tumors rapidly progress
▪ IFNγ and CD40 signaling are nonredundant and critical determinants of intratumoral monocyte fate

## Introduction

Pancreatic ductal adenocarcinoma (PDA) is a lethal malignancy that lacks effective therapies^1^. Therapeutic resistance has been attributed, in part, to the highly suppressive fibroinflammatory tumor microenvironment (TME) characteristic of this disease^2–5^. Tumor-associated macrophages (TAMs) are often the most abundant immune cell infiltrating solid tumors^6^ and can far outnumber tumor cells in PDA^2^. TAMs participate in complex physiologic processes including angiogenesis, metastasis, inflammation and immunity^7^. TAMs with immunosuppressive features correlate with poor PDA patient outcomes^7^, rendering macrophages a desirable therapeutic target.

Pancreatic tumor cells produce numerous pro-myeloid factors such as Ccl2, M-Csf and Gm-Csf thereby promoting myeloid cell expansion and recruitment into the TME^3,5,8–10^. In PDA mouse models that lack a specific tumor antigen, TAMs potently suppress naïve T cell activation^3^. Global TAM abrogation using Csf1r blockade is beneficial in such PDA mouse models^11^. However, clinical efforts to target the CSF1R pathway have not shown therapeutic benefit in PDA patients^12^. Similarly, global TAM depletion using Csf1r blockade fails to promote engineered T-cell antitumor activity in an autochthonous PDA mouse model^13^. TAM depletion shows modest antitumor effects in a poorly immunogenic setting, but not in a parallel animal model in which an endogenous T cell response is engaged^14^. In contrast to TAM depletion, promoting antitumor macrophages using CD40 agonist has shown to be efficacious in murine models and in the clinic^15^. In murine models, CD40 agonist improves the persistence of engineered T cells in PDA^13^ and activity of immune checkpoint blockade^16–18^. However, a recent phase 2 clinical trial with a CD40 agonist + anti-PD1 + chemotherapy failed to improve overall patient survival^19^. Thus, there is a need for a further understanding of macrophage heterogeneity and functionality in PDA to inform novel therapeutic targets.

TAMs are strikingly heterogeneous and can adopt pro-tumorigenic or anti-tumorigenic states that differentially regulate cancer progression. TAMs are derived from the recruitment of circulating monocytes or local proliferation of embryonically seeded tissue resident macrophages^20–22^. Macrophages derived from monocytes display both phenotypic and functional differences compared to tissue resident TAMs, suggesting that ontogeny may partially account for TAM heterogeneity^21,23^. Tissue resident TAMs promote ECM remodeling whereas monocyte-derived macrophages primarily act through shaping immunity^23^. We previously showed engineered T cells induced the accumulation of monocytes in pancreatic tumors that correlated with increased overall survival^24^ and involution of the fibroinflammatory stroma^25,26^. These data support that monocyte-derived macrophages could aid in antigen-specific T cell-mediated destruction of solid tumors. Monocytes are an intriguing therapeutic target because they are continually renewed and expanded in cancer patients^27^ and exhibit plasticity and immunomodulatory capacity. While little is known about the local factors that drive monocyte differentiation into either pro- or anti-tumor states, monocyte differentiation likely hinges on local environmental signals. Notably, T cells co-localize with macrophages in the pancreatic TME^2^ and undergo extensive cross-talk^28^. Together, the data suggest that antigen specific T cells may mediate antitumor effects in part through altering monocytes.

Here, using an in vivo monocyte fate mapping approach, we temporally track monocyte differentiation during tumorigenesis in an orthotopic PDA mouse model that expresses a defined model neoantigen^29^. We identify similar TAM heterogeneity in mouse and human PDA and show most TAMs are derived from recruited monocytes. Further, tumor specific CD4 T cells, but not CD8 T cells, drive monocyte differentiation toward an antitumor TAM state. CD4 T cell depletion leads to exacerbated tumor growth and monocyte-derived TAMs adopting a phenotypic and transcriptional state mirroring immunosuppressive pancreas tissue-resident macrophages. Mechanistically, monocyte acquisition of an antitumor TAM state is dependent on nonredundant IFNγ and CD40 signaling pathways. Loss of these pathways leads to increased tumor growth. Finally, trajectory analyses of PDA patient samples are consistent with a model in which circulating monocytes infiltrating the TME can adopt a protumor or antitumor state, the latter potentially driven by CD40 signaling. Thus, CD4 T cells are a dominant microenvironmental cue governing monocyte fate in cancer. We posit that therapeutic resistance by poorly immunogenic solid tumors may derive, in part, from a de facto monocyte differentiation trajectory toward a pro-tumor state due to a failure to encounter tumor specific CD4 T cells.

## Materials and Methods

### Animals

University of Minnesota Institutional Animal Care and Use Committee approved all animal studies. 6- to 12-wk-old female and male C57BL/6J (000664), *Ifngr1*^-/-^ (003288), *Tnfrsf1a*^-/-^ (003242) and *CX3CR1*^*CreER*^ (B6.129P2(C)-Cx3cr1tm2.1(cre/ERT2)Jung/J) (020940) mice were purchased from The Jackson Laboratory. Sm1x*Rag1-/-* mice^22^ were kindly provided by Dr. Marc Jenkins (University of Minnesota). *CCR2*^creERT2^ [C57BL/6NTac-Ccr2^tm2982^(T2A-Cre7ESR1-T2A-mKate2)] reporter mice^30^ were kindly provided by Burkhart Becker (University of Zurich) were crossed to R26-tdTomato reporter mice (B6.Cg-Gt(ROSA)26Sor^tm9(CAG-tdTomato)Hze^/J). Animals were maintained in SPF facilities maintained by the Research Animals Resources facility at the University of Minnesota where they had free access to food and water and were kept on a 12 hour light-dark cycle.

### Tumor cell lines

The *KPC*2a cell line transduced to express click beetle red luciferase linked to eGFP (CB-eGFP) has been previously described^29^. Tumor cells were cultured in Basic media: DMEM (Life Technologies) + 10% FBS (Life Technologies) + 2.5 mg/ml amphotericin B (Life Technologies) + 100 mg/ml penicillin/streptomycin (Life Technologies) + 2.5 mg dextrose (Fisher Chemical) at 37°C + 5% CO_2_. Medium was sterile filtered and stored in the dark at 4°C. Cell lines used for experiments were maintained below passage 15 and 0.25% trypsin-EDTA (Thermo Fisher) was used to passage tumor cells.

### Orthotopic tumor cell implantation

For orthotopic tumor implantation, mice received 1 mg/kg slow-release buprenorphine injected subcutaneously 2 h prior to surgery for analgesia. Mice were anesthetized using continuous flow of 2-5% isoflurane. Hair was removed using clippers and Nair (Church & Dwight Co., Inc.) and the abdomen was sterilized using a series of 100% EtOH and Betadine washes. Once mice reached surgical plane anesthesia, a small incision was made in the abdomen followed by a small incision in the peritoneum to access the pancreas. 1×10^5^ *KPC2a* cells in 20 μl of 60% Matrigel (Discovery Labware) were injected into the pancreas using an insulin syringe (Covidien) as described^29^. Sutures were used to close the peritoneum (Ethicon) and skin was closed using wound clips (CellPoint Scientific). Mice were monitored daily for 5 days to ensure healing of outer skin.

### Tamoxifen administration

For fate mapping studies, Ccr2 reporter mice were gavaged orally with 250 μl of Tamoxifen (Sigma Aldrich Cat: T5648) at 20 mg ml −1 in corn oil on the day of tumor implantation or 1 day prior to tumor implantation or as indicated.

### In vivo antibody treatments

For T-cell depletion studies, mice were injected intraperitoneally (I.P.) with 200 μg of either anti-CD4 (BioXcell, Cat#-BP0003-1) or anti-CD8 (BioXcell, Cat#-BP0061) on days -1, +2 and +10 post tumor implantation. For CD40L blockade experiments, 500 μg of anti-CD40L (BioXcell, Cat#-BP0017-1) was injected I.P. on days -1 and +2 post tumor implantation.

### Preparation of mononuclear cells from tissues

Spleens were mechanically dissociated to single cells followed by RBC lysis in 1 ml of Tris-ammonium chloride (ACK) lysis buffer (Life Technologies) for 2 min at room temperature (rt). RBC lysis was quenched by addition of 9 ml of T cell media. Splenocytes were centrifuged at 1400 rpm for 5 min, resuspended in T cell media (DMEM, 10% FBS, 2 μM L-glutamine, 100 U/ml penicillin/streptomycin, 25μM 2-β-mercaptoethanol) and kept on ice until further analysis. Tumors were collagenase digested at 37°C for 15 minutes then mechanically digested to single cell suspensions and washed twice to remove cell debris and pancreatic enzymes.

### Cell surface staining

Cells were stained in the presence of 1:500 Fc block (CD16/32, Tonbo) and antibodies diluted 1:200 in FACs buffer (PBS+2.5% FBS) for 45 minutes in the dark at 4°C. Ghost viability dye BV540 (Tonbo) was used to exclude dead cells at 1:500. Cells were fixed in 2% PFA or fixation buffer (Tonbo) for 10-15 at room temperature in the dark prior to data acquisition. Cells were acquired within 24 h using a Cytek Aurora.

| Target Antigen | Conjugate | Reactivity | Clone | Company | Catalog # |
| --- | --- | --- | --- | --- | --- |
| CD45 | BV480 | Mouse | 30-F11 | BD | 566168 |
| Ly6C | BV421 | Mouse | HK1.4 | Biolegend | 128031 |
| CD11b | BV605 | Mouse | M1/70 | Biolegend | 101237 |
| F4/80 | BV650 | Mouse | BM8 | Biolegend | 123149 |
| CD64 | BV711 | Mouse | 10.1 | Biolegend | 305041 |
| Ly6G | BV785 | Mouse | 1A8 | Biolegend | 127645 |
| CD206 | PerCP Cy5.5 | Mouse | C068C2 | Biolegend | 141715 |
| PDL1 | PeCy7 | Mouse | 10F.9G2 | Biolegend | 124313 |
| FRβ | APC | Mouse | 10/FR2 | Biolegend | 153305 |
| Arg-1 | PeCy7 | Mouse | A1exF5 | ThermoFisher | 25369782 |
| MHCII | APC Fire 750 | Mouse | M5/114.15.2 | Biolegend | 107651 |
| Ki67 | Alexa Flour 700 | Mouse | 16A6 | Biolegend | 652420 |
| CD3 | BV650 | Mouse | 17A2 | BD | 740530 |
| CD8 | BV711 | Mouse | 53-6.7 | Biolegend | 100759 |
| CD44 | PerCP Cy5.5 | Mouse | IM7 | Tonbo | 650042U100 |
| CD4 | AF700 | Mouse | RM4-5 | ThermoFisher | 56004282 |
| Klrg1 | PeCy7 | Mouse | 2F1 | eBioscience | 25589382 |
| Foxp3 | FITC | Mouse | FJK-16s | eBioscience | 48577382 |
| T-bet | APC | Mouse | 4B10 | Biolegend | 644810 |
| Stat 1 | PE | Mouse | D1K9Y | Cell Signaling | 80916S |
| IFNγ | PeCy7 | Mouse | XMG1.2 | Invitrogen | 24731141 |
| CD19 | FITC | Mouse | 1D3 | Biolegend | 152403 |
| CD115 | PerCP Cy5.5 | Mouse | AFS98 | Biolegend | 135525 |
| Lyve1 | ef660 | Mouse | ALY7 | ThermoFisher | 50-443-82 |
| CD11b | APC | Human | ICRF44 | Biolegend | 301310 |
| CD11c | PeCy7 | Human | 3.9 | eBioscience | 25016642 |
| CD64 | APC Fire 750 | Human | 10.1 | Biolegend | 305036 |
| CD14 | FITC | Human | 61D3 | ThermoFisher | 11014942 |
| CD15 | BV650 | Human | HI98 | BD | 564232 |
| FRβ | PE | Human | 94b | Biolegend | 391703 |
| HLA-DR | PerCP Cy5.5 | Human | L243 | BD | 552764 |
| PD-L1 | BV711 | Human | 29E.2A3 | Biolegend | 329722 |
| CD163 | BV785 | Human | GHI/61 | Biolegend | 333632 |

### Intracellular staining

The Foxp3 intracellular staining kit (Tonbo) was used for detecting intracellular transcription factors and proteins. Following cell surface staining, cells were washed 2X in FACs buffer, fixed for 30 min at 4°C, washed 2X in permeabilization buffer, stained with antibodies diluted 1:100 in permeabilization buffer for 1-2 hours in the dark at 4°C. Cells were washed 2X in permeabilization buffer, resuspended in FACs buffer and acquired within 24 h on a Cytek Aurora flow cytometer following addition of cell counting beads (Sigma).

### PMA/Ionomycin restimulation

To determine ex vivo T cell functionality from tumor-bearing mice, single cell suspensions from spleen and tumor were obtained as described above and activated *in vitro* as described^14,31^. Briefly, mononuclear cells were restimulated with 1X Cell Stimulation Cocktail (eBioscience) in the presence of Golgiplug and Golgistop (BD) according to manufacturer’s instructions in T cell media. 4-5 hours later, cells were stained with live/dead ghost dye at 1:500 (Tonbo) and the following antibodies at diluted in FACs buffer at 1:200 against CD45 (30F-11, BD), CD3 (17A2, Biolegend), CD4 (RM4.5, Tonbo), CD8α (53-6.7, Tonbo), Klrg1 (2F1, eBioscience), and CD44 (IM7, Tonbo) for 30 minutes at 4°C in the dark.

Cells were washed 2X in FACs buffer, fixed/permeabalized using the BD cytofix/cytoperm kit (BD), and stained with anti-IFNγ (XMG1.2, Biolegend) diluted 1:100 in perm/wash buffer for 1 h at 4°C. Cells were washed 2X in perm/wash buffer, resuspended in FACs buffer and stored overnight at 4°C in the dark. Cells were acquired the following day on a Fortessa 1770 flow cytometer following the addition of counting beads (Sigma) and analyzed using FlowJo software (version 10).

### Murine tumor scRNAseq sample acquisition and data analysis

For scRNAseq, 4 tumors from each group were harvested and processed to generate single cell suspensions. Live CD45+ Tomato+ cells and CD45+ Tomato-cells were FACS sorted using a BD FACSAria II. Each population was then stained with hashtag oligo antibodies (BD Biosciences; HTO#9 and HTO#10) and BD Bioscience CITEseq antibodies: CD274 (Cat:153604), IA-IE (Cat:107653), CD11b (Cat: 101265), Folate receptor beta (Cat: 153307) for 30 mins at 4°C (1:500) then recombined at a 1:1 mix. Sorted cells were resuspended in a final concentration of 100 cells per μl in 1X PBS containing 0.04% BSA for single cell capture of approximately 20,000 cells per group. Cells were submitted to University of Minnesota Genomics Core (UMGC) for single cell 10X Chromium 3’ GEX Capture and NovaSeq 2 x150 S4 sequencing targeting ∼50,000 reads per cell.

For preprocessing of the mouse scRNAseq data, we removed genes detected in less than 10 cells, potential empty cells with less than 200 feature counts, and apoptotic cells possessing more than 25% mitochondrial mRNA content. We then utilized DoubletFinder^32^ to perform a more elegant doublet removal independently for each sequence capture prior to data merging and integration. NormalizeData and ScaleData functions from Seurat (v4.0.1) were used for normalization and scaling. Variable features were extracted using FindVaribleFeatures function. For integration purpose, we chose Harmony package^33^. The first 20 principle components were used to generate uniform manifold approximation and projection (UMAP) and t-distributed stochastic neighbor embedding (tSNE). For gene set scores, Seurat AddModuleScore function was used, where pro-inflammatory (*Il1a, Cxcl12, Il1b, Cxcl3, Tnf*) and anti-inflammatory (*Mrc1, Il10, Siglec1, FRβ, Arg1*) gene sets were used. SingleR (v1.6.1)^34^ was used as an unbiased computational method for immune cell type annotation. Clusters generated using resolution 0.1 that were identified as SingleR monocyte and macrophage were extracted for further sub-clustering, pseudo-time trajectory and intercellular interaction analysis. Trajectory analysis were performed with support of Monocle3 (v1.0.0)^35^. In order to avoid batch effects across captures, we applied “harmony” as the base reduction method for all Monocle3 functions. Cells in the monocyte cluster were used as the root population to calculate pseudo-time inference of monocyte-macrophage differentiation. Pseudo-time parameter from ordered cells was extracted for visualization in “harmony” embeddings. For intercellular communication analysis, we utilized NicheNet (v1.0.0)^36^. All NicheNet models were first converted to mouse gene symbols using convert_human_to_mouse_symbols function.

### Bulk RNA sequencing collection and analysis

Tumor single cell suspensions were isolated from 4 WT and 4 *Ifngr1*-/- mice and CD45+ CD11b+ F4/80+ MHCII^hi^ and MHCII^lo^ macrophages were FACS sorted into Trizol for RNA extraction. A minimum of 10,000 cells were sorted and submitted to UMGC for RNA isolation and sequencing using the Novaseq platform. Bulk RNAseq processing was performed using CHURP pipeline developed by the Minnesota Supercomputing Institute, which implemented and integrated Trimmomatic, HISAT2, SAMTools and featureCounts. Mus musculus GRCm38 (Ensembl release 102) was used as mouse reference genome. Differential expression analysis was adopted from DEseq2 (v.1.32.0). Pathway analyses were performed using fgsea function from the fgsea package (v.1.18.0).

### Human PDA samples

De-identified and resected human tumor and normal adjacent tissues were obtained from BioNET, a University of Minnesota IRB-approved protocol and tissue bank for investigators. Tumors were from patients diagnosed with PDA. Single cell mononuclear suspensions from tissues specimens were generated like mouse tumors as described above. Cells were stored at -80°C in Cryostore and thawed for staining and flow cytometric analysis like mouse samples.

### Human PDA scRNAseq data analysis

Publicly available scRNAseq data from tumors from 6 PDA patients available from Elyada et al^40^ was downloaded after NIH approval at dbGaP (accession number phs001840.v1.p1). Filtered count matrices for 6 human tumor samples (SRR9274536, SRR9274537, SRR9274538, SRR9274539, SRR9274542, SRR9274544) were used as input data. Analytic tools used for human PDA scRNAseq data was identical to that of mouse scRNAseq described above, with the exception that SingleR model training was performed using HumanPrimaryCellAtlasData from Celldex.

### Statistical analysis

Statistical analyses were performed using GraphPad software (version 9.0). Mouse experiments include n=3-8 mice per group. Unpaired, two-tailed Student’s T test was used to compare two-group data. One-way ANOVA and Tukey posttest were used for comparing >2-group data. Data are presented as mean ± standard error of the mean (SEM), and *p*<0.05 was considered significant. \**p*<0.05, \*\**p*<0.005, and \*\*\**p*<0.0005.

## Results

### Fate mapping of monocyte differentiation in PDA

Given that both monocyte-derived and embryonically-derived macrophages accumulate in PDA, distinguishing monocyte derived macrophages in vivo has been challenging. To overcome this, we utilized the CCR2^CreER^ R26^Tdtomato^ monocyte fate mapping mouse that allows for specific labeling of individual waves of blood monocytes following tamoxifen treatment, enabling tracking of their differentiation upon entry into tissue^37–39^. Tamoxifen treatment of CCR2^CreER^ R26^Tdtomato^ mice in steady state revealed robust classical monocyte labeling in the blood, but no labeling of pancreatic tissue resident macrophages (Supplemental figure 1A-D), consistent with their reported lack of Ccr2 expression^40^. To track monocyte differentiation in PDA in the presence of tumor specific T cells, we orthotopically implanted *KPC2a* tumor cells that expresses a model neoantigen click beetle red luciferase (CB)^29^ into the pancreas of CCR2^CreER^ R26^Tdtomato^ mice following a single dose of tamoxifen (Figure 1A). We analyzed mice at day 3, 7, and 14 after implantation and identified two distinct Tomato+ monocyte derived macrophage subsets based on MHCII^hi^ and MHCII^lo^ phenotype (Figure 1B). MHCII^hi^ macrophages expressed higher levels of CD86, consistent with a more immunostimulatory phenotype, whereas MHCII^lo^ macrophages expressed markers of alternative activation like Arg-1 (Figure 1C). Moreover, intratumoral Tomato+ macrophages increased from day 3 to day 7, then decreased by both percentage and number by day 14 (Figure 1D), which is likely due to replacement by subsequent waves of non-labeled recruited monocytes. On day 3 post tumor and tamoxifen, most Tomato+ cells resembled undifferentiated monocytes (CD11b+, Ly6C+, F4/80-). However, by day 7 and 14, most Tomato+ cells expressed markers of fully differentiated macrophages (Figure 1E), supporting that recruited monocytes differentiate into macrophages as early as day 7 and are maintained for at least an additional week. At day 3, Tomato+ macrophages (CD64+F480+Ly6C-) were MHCII^lo^ whereas by days 7 and 14, most Tomato+ macrophages were MHCII^hi^. Thus, monocytes undergo an initial differentiation into a transitory MHCII^lo^ state that subsequently become MHCII^hi^. Alternatively, MHCII^hi^ monocyte-derived macrophages may preferentially expand and/or survive during tumor progression (Figure 1F).

**Figure 1.**
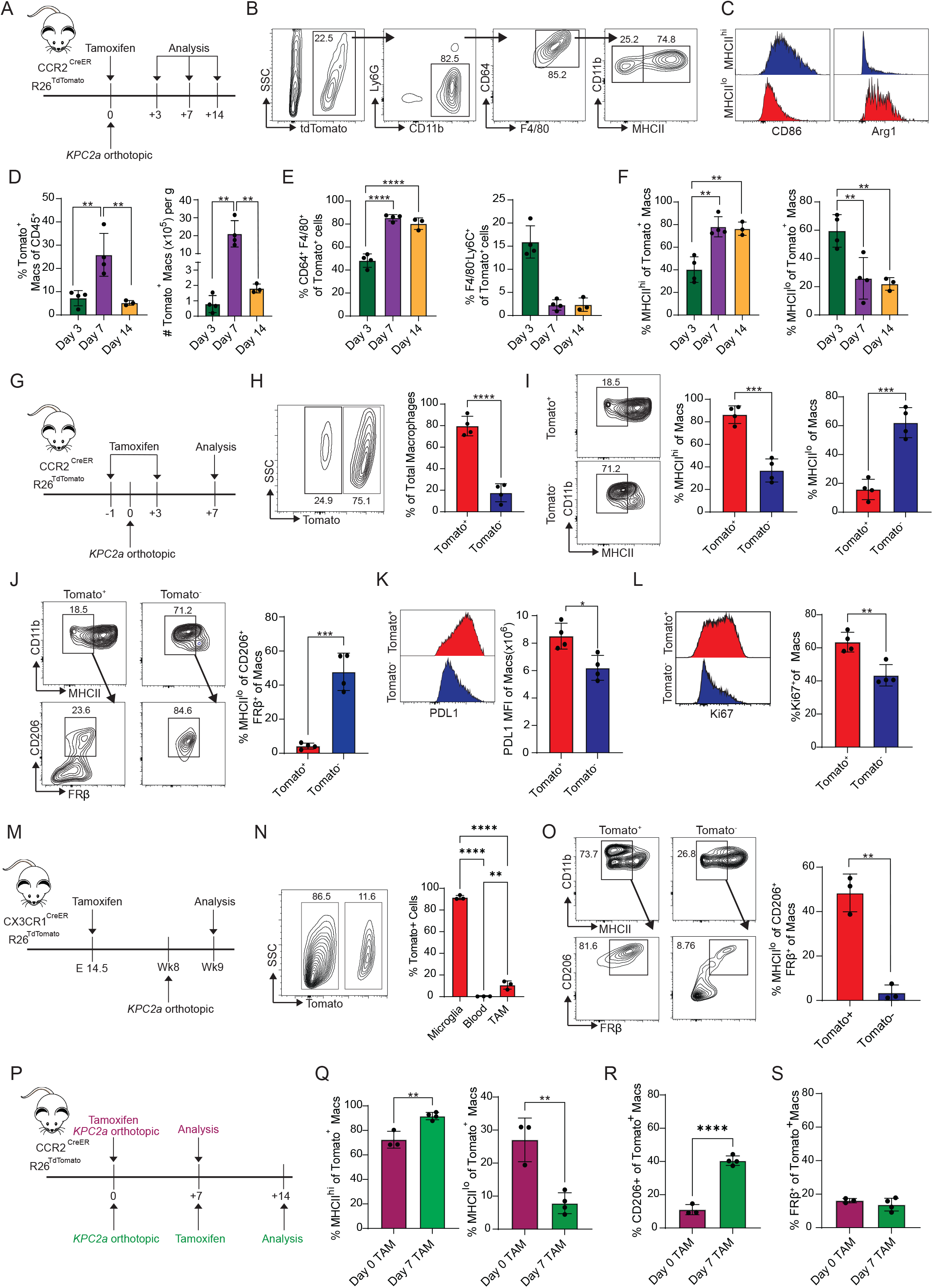
Fate Mapping of Monocyte Differentiation in PDA. A) *In vivo* monocyte tracking approach in CCR2^CreER^ R26^TdTomato^ mice orthotopically implanted with *KPC2a* tumor cells and treated with tamoxifen on the same day of tumor implantation. B) Gating strategy for distinguishing Tomato+ MHCII^hi^ and MHCII^lo^ macrophage subpopulations. C) Representative CD86 and Arg1 histograms gated on Tomato+ MHCII^hi^ or MHCII^lo^ intratumoral macrophages on day 7. D) Frequency and number of Tomato+ TAMs. Each dot is an independent mouse. Data are mean ± S.E.M. n=3-4 mice per group. \*\**p*<0.005. One-way ANOVA with a Tukey’s posttest. E) Proportion of intratumoral Tomato+ cells that are macrophages (CD64+ F4/80+) or monocytes (F4/80-Ly6C+). Each dot is an independent mouse. Data are mean ± S.E.M. n=3-4 mice per group. \*\*\*\**p*<0.0001. One-way ANOVA with a Tukey’s posttest. F) Proportion of intratumoral Tomato+ macrophages that are MHCII^hi^ or MHCII^lo^. Each dot is an independent mouse. Data are mean ± S.E.M. n=3-4 mice per group. \*\**p*<0.005. One-way ANOVA with a Tukey’s posttest. G) *In vivo* monocyte tracking approach in CCR2^CreER^ R26^TdTomato^ mice orthotopically implanted with *KPC2a* tumor cells and treated with tamoxifen twice for continuous labeling. H) Representative plot gated on intratumoral F4/80+CD64+ macrophages on day 7. Proportion of total macrophages that are Tomato+. Each dot is an independent mouse. Data are mean ± S.E.M. n=4 mice per group. \*\*\*\**p*<0.0001, Student’s t-test. I) Representative plots gated on day 7 and proportion of Tomato+ or Tomato-intratumoral MHCII^hi^ or MHCII^lo^ macrophages on day 7. Each dot is an independent mouse. Data are mean ± S.E.M. n=4 mice per group, \*\*\**p*<0.001, Student’s t-test. J) Representative plots gated on day 7 and proportion of Tomato+ and Tomato-MHCII^lo^CD206+FRβ+ macrophages on day 7. Each dot is an independent mouse. Data are mean ± S.E.M. n=4 mice per group. \*\*\**p*<0.0001, Student’s t-test. K) Representative histograms and PD-L1 mean fluorescence intensity (MFI) of Tomato+ and Tomato-macrophages at day 7. Each dot is an independent mouse. Data are mean ± S.E.M. n=4 mice per group. \*\*\**p*<0.0001, Student’s t-test. L) Representative histograms and frequency of Tomato+ or Tomato-macrophages that are Ki67+ on day 7. Each dot is an independent mouse. Data are mean ± S.E.M. n=4 mice per group. \*\*\**p*<0.0001, Student’s t-test. M) *In vivo* embryonic fate mapping approach in CX3CR1^CreER^ R26^TdTomato^. Briefly, pregnant mothers were gavaged with tamoxifen on embryonic day 14.5 then progeny were implanted with tumors and assessed 7 days later. N) Representative plot gated on intratumoral F4/80+CD64+ macrophages on day 7 from mice in M. Proportion of microglia, CD45+ blood immune cells and tumor macrophages that are Tomato+. Each dot is an independent mouse. Data are mean ± S.E.M. n=3 mice per group. \*\**p*<0.005, \*\*\*\**p*<0.0001, Student’s t-test. O) Representative plots gated on day 7 and proportion of Tomato+ and Tomato-MHCII^lo^CD206+FRβ+ tumor macrophages on day 7 from mice in M. Each dot is an independent mouse. Data are mean ± S.E.M. n=3 mice per group. \*\**p*<0.005, Student’s t-test. P) Experimental schematic to to label the initial wave of recruited monocytes (purple font) or subsequent waves of recruited monocytes (green font). Q) Proportion of each monocyte wave that gives rise to MHCII^hi^ or MHCII^lo^ macrophages. Each dot is an independent mouse. Data are mean ± S.E.M. n=3-4 mice per group. \*\**p*<0.005, Student’s t-test. R) Proportion of each monocyte wave that gives rise to CD206+ macrophages. Each dot is an independent mouse. Data are mean ± S.E.M. n=3-4 mice per group. \*\*\*\**p*<0.0001, Student’s t-test. S) Proportion of each monocyte wave that gives rise to FRβ+ macrophages. Each dot is an independent mouse. Data are mean ± S.E.M. n=3-4 mice per group.

We next compared the phenotype of Tomato+ monocyte-derived macrophages to Tomato-tissue resident macrophages. We administered a tamoxifen at day -1- and 3-days following tumor implantation (Figure 1G), which resulted in nearly 100% monocyte labeling for 7 days (Supplemental figure 1E). Nearly 80% of the tumor infiltrating macrophages were Tomato+ (Figure 1H), again supporting that TAM pool is mostly monocyte derived. Consistent with prior reports^23^, non-monocyte derived TAMs primarily adopted an MHCII^lo^ phenotype (Figure 1I). Further, MHCII^lo^ Tomato-macrophages also expressed markers of suppressive macrophages including CD206 and FRβ. In contrast, Tomato+ macrophages rarely adopted a MHCII^lo^ CD206^+^ FRβ^+^ phenotype (Figure 1J) and instead expressed high PD-L1 (Figure 1K) and were more proliferative (Figure 1L). These data highlight immune modulatory and proliferation differences between macrophage subpopulations. Macrophages in the non-tumor-bearing healthy pancreas exhibited an MHCII^lo^CD206^+^ FRβ^+^ phenotype, supporting that this population is derived from tissue resident macrophages (Supplemental figure 2). To investigate the developmental origin of pancreatic tissue resident macrophages in PDA, we performed embryonic pulse chase experiments in tumor bearing CX3CR1^CreER^ R26^TdTomato^ mice. Given that CX3CR1 is turned on in embryonic derived macrophages during development^41^, in utero treatment of tamoxifen allows for robust and specific labeling of embryonic macrophages in these mice^38^. Pregnant CX3CR1^CreER^ R26^TdTomato^ mice were administered tamoxifen on embryonic day 14.5, then their progeny were implanted with tumors at 8 weeks of age then assessed 7 days post tumor (Figure 1M). Nearly 100% of microglia were tomato+ while labeling in the blood was negligible (Figure 1N). Approximately 10% of the tumor macrophages were Tomato+ (Figure 1N), confirming that the vast majority of TAMs are deriving from non-tissue resident macrophages. Furthermore, Tomato+ cells primarily adopted an MHCII^lo^CD206^+^ FRβ^+^ phenotype while Tomato-cells rarely did (Figure 1O), supporting that MHCII^lo^CD206^+^ FRβ^+^ TAMs are derived from embryonic tissue resident macrophages.

As tumor burden could impact monocyte differentiation, we next compared monocyte-derived macrophage fate during early tumorigenesis to those that are recruited later. We administered tamoxifen to label monocytes on day 0 (early) or day 7 (late) following tumor implantation and assessed monocyte-derived macrophage phenotype 7 days following tamoxifen (Figure 1P). Monocytes recruited later in tumor development preferentially differentiated into MHCII^hi^ macrophages compared to initially recruited monocytes (Figure 1Q). For monocytes infiltrating in larger tumors, a subset did upregulate CD206 (Figure 1R) suggesting a potential acquisition of a suppressive phenotype. At both timepoints, Tomato+ macrophages failed to upregulate FRβ (Figure 1S) suggesting FRβ is preferentially expressed by pancreas resident macrophages.

### CD4 depletion promotes pro-tumor macrophage differentiation

T cells co-localize with intratumoral macrophages and interact with both monocytes and TAMs^2^. Given these observations, we hypothesized that the presence of a tumor-specific T cell response may be a key determinate of monocyte differentiation. To test this concept, we depleted CD4 or CD8 T cells in CCR2^creER^ R26^tdTomato^ mice bearing tumors and assessed monocyte differentiation at day 7 (Figure 2A). CD8 T cell depletion led to a minor expansion of Tomato+ macrophages, suggesting a role for CD8 T cells in regulating monocyte recruitment, differentiation and/or survival (Figure 2B). While CD8 T cell depletion had minimal effect on Tomato+ macrophage phenotype, CD4 T cell depletion led to a dramatic reduction of MHCII^hi^ macrophages and a corresponding expansion of the MHCII^lo^ population (Figure 2C). These data support that CD4 T cells promote monocyte differentiation into MHCII^hi^ macrophages. To test if CD4 T cells were merely modulating MHCII, or instead shifting macrophage subsets, we developed an expanded antibody panel to delineate macrophage subpopulations in CD4 T cell depleted and control tumor-bearing mice at days 7 and 14 after tamoxifen (Figure 2D). At day 14, tumor weights were significantly increased in CD4 T cell depleted as compared to control mice demonstrating a critical role for CD4 T cells in limiting tumor growth (Figure 2E). Moreover, CD4 T cell depletion again resulted in a marked expansion of MHClI^Io^ macrophages at day 14 compared to untreated controls (Figure 2F). At both time points, a significantly higher proportion of Tomato+ macrophages expressed pro-tumorigenic markers including Arg-1, CD206 and FRβ in CD4-depleted vs. control mice (Figure 2G-H), supporting a monocyte differentiation program switch when CD4 T cells are absent. Furthermore, CD4 T cell depletion led to a downregulation of PD-L1 on Tomato+ macrophages (Figure 2I) and nearly half of the Tomato+ macrophages adopted a MHCII^lo^ CD206+FRβ+ tissue-resident phenotype that is largely absent in control mice (Figure 1J). Finally, CD4 T cell depletion led to a dramatic increase in the proliferation of non-monocyte-derived tissue resident macrophages (Figure 2K). These data suggest that CD4 T cells limit monocyte differentiation and/or proliferation toward a pro-tumorigenic tissue resident-like state. Overall, these studies support that CD4 T cells are a key determinate of monocyte fate toward an MHCII^hi^ antitumor state and restrain their ability to differentiate into suppressive Arg1+ TAMs and CD206^+^FRβ^+^ TAMs, the latter indistinguishable from tissue resident TAMs.

**Figure 2.**
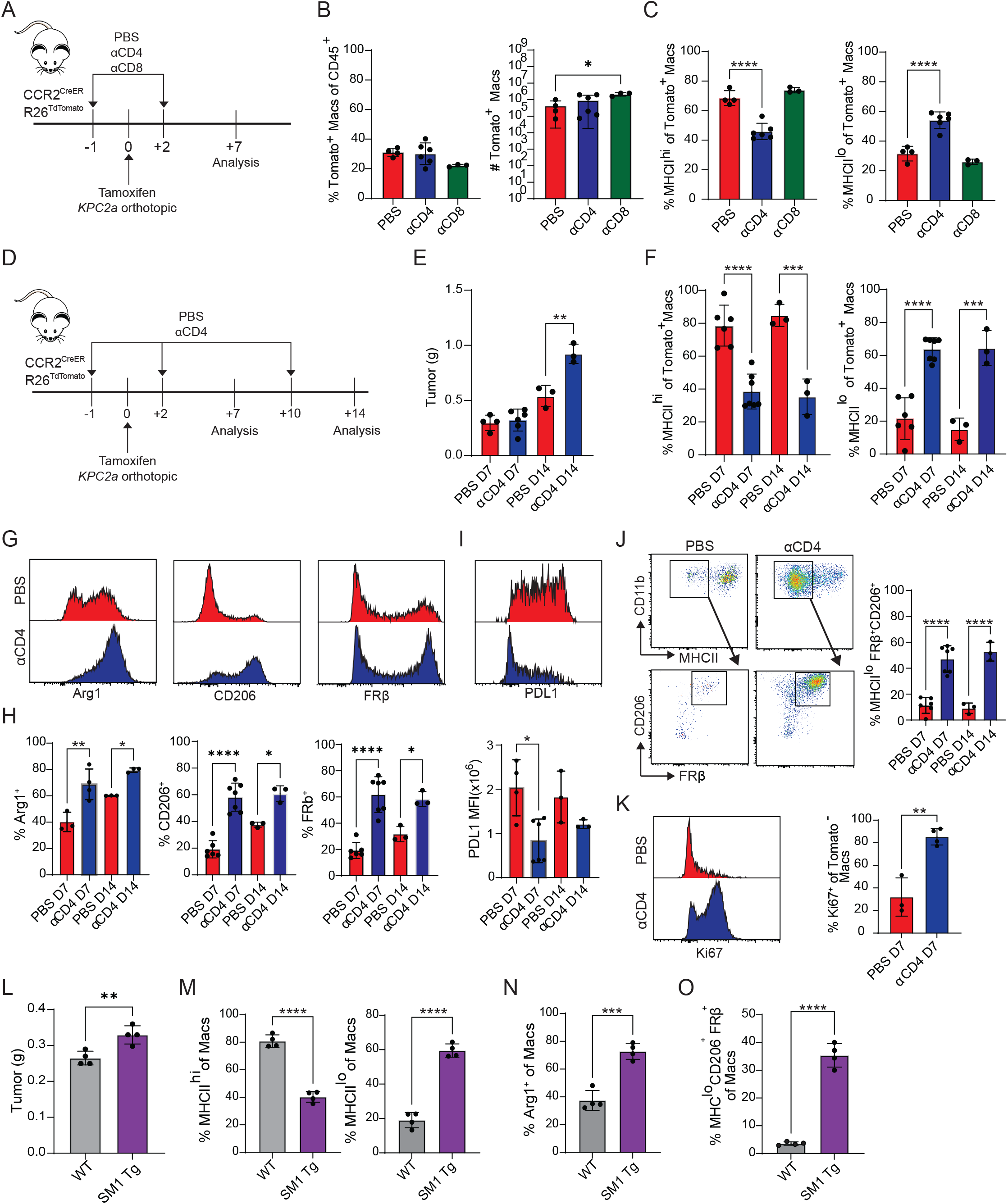
Tumor specific CD4 T cells drive MHCII^hi^ macrophage differentiation in PDA. A) Experimental approach to test the role of CD4 and CD8 T cells on monocyte differentiation in PDA. B) Proportion (left) and number (right) of Tomato+ intratumoral macrophages gated on CD45+ cells. Each dot is an independent mouse. Data are mean ± S.E.M. n=3-5 mice per group. \**p*<0.05. One-way ANOVA with a Tukey’s posttest. C) Proportion of Tomato+ macrophages that are MHCII^hi^ or MHCII^lo^. Each dot is an independent mouse. Data are mean ± S.E.M. n=3-5 mice per group. \*\*\*\**p*<0.0001. One-way ANOVA with a Tukey’s posttest. D) Experimental approach to test the impact of prolonged CD4 T cell depletion in tumor bearing CCR2_CreER_ R26_TdTomato_ mice. E) Tumor weight in grams (g) from mice in D. n=3-6 mice per group. Data are mean ± S.E.M. \*\**p*<0.005. One-way ANOVA with a Tukey’s posttest. F) Proportion of Tomato+ macrophages that are MHCII^hi^ or MHCII^lo^ from mice in E. Data are mean ± S.E.M. n=3-6 mice per group. \*\*\**p*<0.001, \*\*\*\**p*<0.0001. One-way ANOVA with a Tukey’s posttest. G) Representative histograms of Arg1, CD206, and FRβ gated on intratumoral Tomato+ macrophages isolated from saline or αCD4 treated mice. H) Proportion of intratumoral Tomato+ macrophages expressing Arg1, CD206, and FRβ from saline or αCD4 treated mice. Data are mean ± S.E.M. n=3-6 mice per group. \**p*<0.05, \*\**p*<0.005, \*\*\*\**p* <0.0001, separate Student’s t-test for each timepoint. I) Representative PDL1 staining (top) and MFI (bottom) by intratumoral Tomato+ macrophages from saline or αCD4 treated mice. n=3-6 mice per group. \**p*<0.05, Student’s t-test. J)Representative plots and proportion of Tomato+ MHCII^lo^ macrophages that co-express CD206 and FRβ from mice in E. Data are mean ± S.E.M. n=3-6 mice per group. \*\*\*\**p*<0.0001, Student’s t-test. K) Representative histograms and proportion of Tomato-macrophages that are Ki67+. Data are mean ± S.E.M. n=3-4 mice per group. \*\**p*<0.005, Student’s t-test. L) Tumor weight in grams (g) from wild type (WT) or SM1xRAGKO mice 7 days after *KPC2a* implantation. Data are mean ± S.E.M. n=4 mice per group. \*\**p*<0.005, Student’s t-test. M) Proportion of MHCII^hi^ or MHCII^lo^ macrophages from mice in L. Data are mean ± S.E.M. n=4 mice per group. \*\*\*\**p*<0.0001, Student’s t-test. N) Proportion of Arg1+ macrophages + from mice in L. Data are mean ± S.E.M. n=4 mice per group. \*\*\**p*<0.001, Student’s t-test. O) Proportion of MHCII^lo^CD206+FRβ+ macrophages from mice in L. Data are mean ± S.E.M. n=4 mice per group. \*\*\*\**p*<0.0001, Student’s t-test.

### Tumor specific CD4 T cells drive MHCII^hi^ macrophage differentiation

We next tested if tumor antigen-specific CD4 T cell were responsible for promoting MHCII^hi^ TAMs by implanting *KPC2a* cells orthotopically into wild type (WT) or CD4 SM1 TCR transgenic *Rag1*-/- mice, in which CD4 T cells express a monoclonal TCR specific to an irrelevant Salmonella antigen^42^. Using this approach, any potential endogenous tumor specific CD4 T cell is eliminated. *KPC*2a tumors in SM1 mice were significantly larger as early as day 7 (Figure 2L) supporting a role for tumor specific CD4 T cells in controlling tumor growth. MHCII^lo^ TAM frequency was markedly increased in SM1 mice (Figure 2M). Furthermore, Arg1+ (Figure 2N) and MHC^lo^CD206^+^FRβ^+^ TAMs (Figure 2O) were also increased in SM1 mice. While intratumoral activated CD44^hi^ CD4 T cell frequency was reduced in SM1 mice, CD4 T cell number were unchanged (Supplemental figure 3), demonstrating that bystander/non-specific CD4 T cells still infiltrate PDA. Taken together, tumor specific CD4 T cells govern the program of monocyte differentiation into MHCII^hi^ antitumor TAMs and in their absence, monocytes default to distinctly suppressive phenotypes.

### Single Cell Fate Mapping of Monocyte differentiation in PDA

We next sought to map monocyte differentiation trajectories in the presence or absence of CD4 T cells. As such, we performed single cell RNA and CITE sequencing (scRNAseq) of sorted CD45+ Tomato+ and CD45+ Tomato-immune cells from orthotopic tumors from control or CD4-depleted CCR2^CreER^ R26^Td tomato^ mice. Control tumors were harvested at day 3 and day 7, and CD4-depleted tumors were harvested at day 7 post monocyte labeling and tumor implantation. Clustering of integrated data at the 2 time points using Seurat revealed a diversity of immune cell populations residing within tumors including monocytes, macrophages, T cells, B cells, and dendritic cells (Figure 3A and Supplemental figure 4). Sub-clustering of monocyte/macrophage populations revealed 5 populations (Figure 3B). Cluster 1 expressed high levels of monocyte specific genes including *Ly6c2, Ms4a4c* and *Plac8* (Figure 3C). Clusters 2, 3 and 4 expressed macrophage specific genes such as *Adgre1, Cd68* and *Fcgr1*, suggesting 3 distinct macrophage populations (Supplemental figure 5). Cluster 3 expressed *H2-Ab1* (MHC II) and other genes associated with antigen presentation (*H2-Aa, Cd74*), while Cluster 4 expressed immunosuppressive genes like *Arg1* and *Spp1* (Figure 3C), mirroring the MHCII^hi^ or MHCII^lo^ Arg-1+ populations (Figure 1B). Both Arg1^43^ and Spp1^44^ have immunosuppressive properties, suggesting a pro-tumorigenic functions of Cluster 4. Cluster 2 consisted of cells enriched for *Folr2, Lyve1*, and *Cd206* (Figure 3C and Supplemental figure 5) that are associated with a tissue resident macrophage phenotype^45,46^. Cluster 5 primarily consisted of proliferating cells, as shown by abundant *Mki67* and *Top2a* transcripts (Figure 3C). Analysis of protein expression using CITE sequencing showed that MHCII expression was almost exclusively restricted to cluster 3, while *Folr2* was primarily expressed by cluster 2 (Figure 3D). Furthermore, PD-L1 was expressed by both cluster 3 and 4 yet was absent from the *Folr2*+ cluster 2, confirming the flow cytometric data (Figure 1). We next compared gene expression in macrophage clusters to canonical pro-inflammatory and anti-inflammatory macrophage gene sets. Cluster 3 (MHCII^hi^) expressed more genes associated with a pro-inflammatory state, including *Cxcl9* and *Cxcl10* that are IFNγ-inducible chemokine ligands promote effector T cell migration, highlighting the potential antitumor role of MHCII^hi^ macrophages. In contrast, cluster 2 (*Folr2*^+^) and cluster 4 (MHCII^lo^ *Arg1*+) were enriched for anti-inflammatory genes such as *Il10* (Figure 3E).

**Figure 3.**
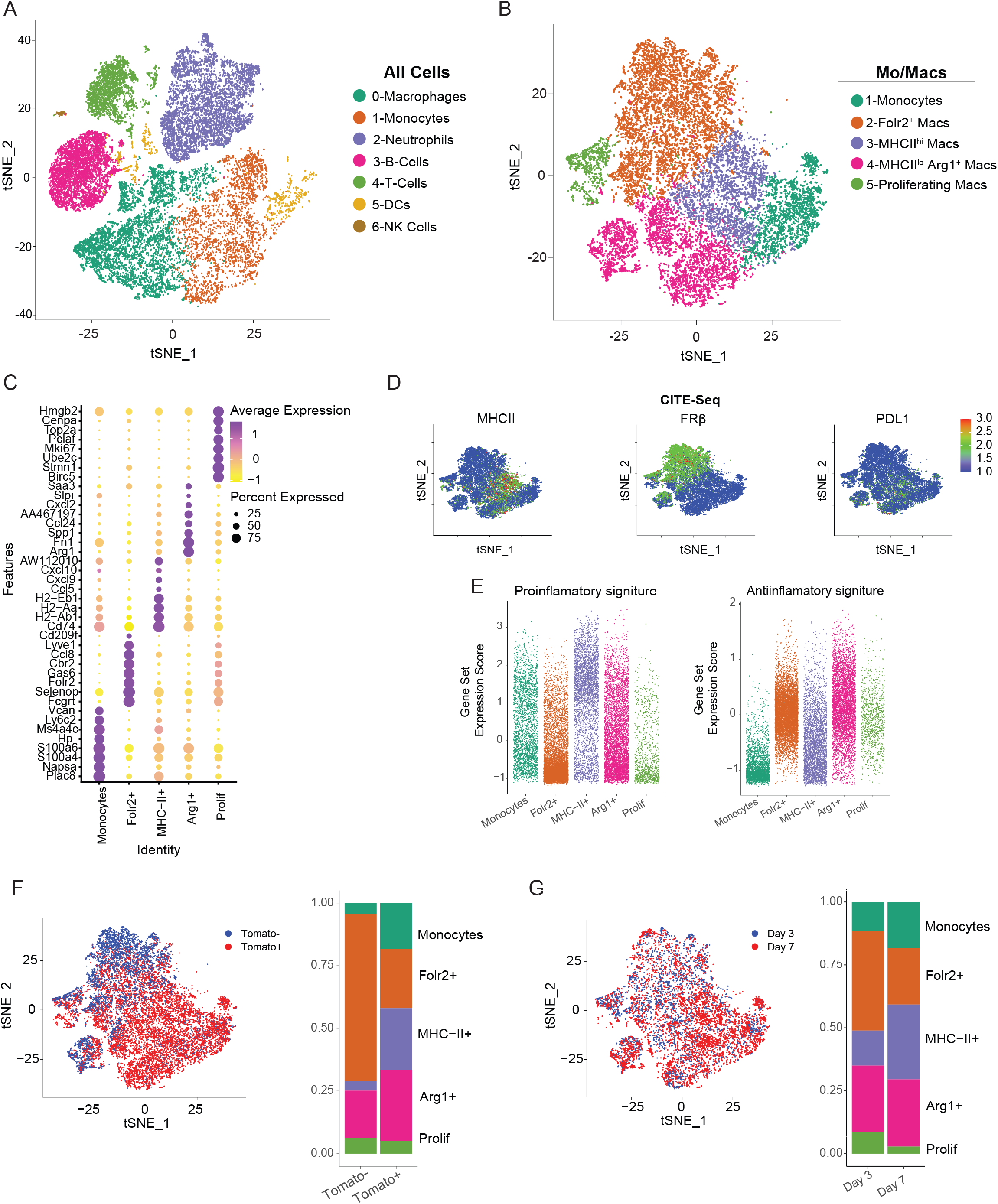
Single Cell fate mapping of monocyte differentiation in PDA. A) t-distributed stochastic neighbor embedding (tSNE) plots of intratumoral immune cells isolated on day 3 and day 7 ± αCD4 posttumor were generated from scRNAseq data. Live, CD45+Tomato+ and live CD45+Tomato-cells were sorted, HTO labeled and recombined for sequencing (n=4 orthotopic *KPC*2a tumors per group). Cell types were named using singleR. B) tSNE plots of only monocytes and macrophage clusters from A. Monocyte and macrophage clusters were selected based on singleR delineation C) Heatmap showing normalized gene expression levels of cluster-specific genes. D) Normalized protein expression of CITEseq antibodies in tSNE plot from B. E) Gene set enrichment of canonical proinflammatory and anti-inflammatory genes split by monocyte/macrophage cluster defined in B. F) tSNE plot showing Tomato+ and Tomato-cells from Day 3 and Day 7 untreated and proportion of each monocyte/macrophage cluster among Tomato+ and Tomato-myeloid cells from clusters defined in B (right). G) tSNE plot and proportion of each monocyte/macrophage cluster at day 3 and day 7 untreated.

We next assessed the relative contribution of Tomato+ and Tomato-cells to the TAM pool and found that most Tomato-macrophages clustered within the *Folr2*+ population, whereas Tomato+ macrophages primarily clustered within the monocyte, MHCII^hi^ and Arg1+ populations (Figure 3F, Supplemental figure 6A). Together, these data support that MHCII^hi^ and Arg1+ macrophages are derived from monocytes, and FRβ+ (*Folr2*+) macrophages are derived from tissue resident macrophages.

To examine temporal changes in macrophage populations, we compared the relative proportion of each cluster at day 3 and day 7. At day 3, most macrophages clustered within the Folr2+ and Arg1+ populations, However, by day 7, macrophages adopting the MHCII^hi^ phenotype expanded (Figure 3G, Supplemental figure 6B), consistent with our flow cytometry data showing the skewing of monocyte differentiation toward MHCII^hi^ over time (Figure 1F). Notably, the day 7 timing of MHCII^hi^ TAM enrichment corresponds to the timing of a peak of an antigen specific CD4 T cell response.

### CD4 T cell depletion rewires monocyte differentiation toward a protumor state

To further assess the role of CD4 T cells on monocyte fate, we clustered Tomato+ cells from CD4 T cell depleted or control tumors. Most Tomato+ cells clustered in the MHCII^hi^ population in tumors from PBS treated mice (Figure 4A, B). In contrast, CD4 T cell depletion caused a dramatic increase in the proportion of Tomato+ cells adopting an Arg1+ or Folr2+ transcriptional profile while a concomitant reduction in Tomato+ cells adopting an MHCII^hi^ transcriptional profile was observed (Figure 4A, B). Furthermore, monocyte derived macrophages upregulated additional genes associated with tissue resident macrophages including *Lyve*1 in CD4 T cell depleted tumors (Supplemental figure 7). *Lyve*1 is previously thought to be restricted to embryonically-derived cells^46^. Thus, in the absence of CD4 T cells, a subset of monocyte-derived macrophages adopt a phenotypic and transcriptional profile mirroring tissue resident macrophages.

**Figure 4.**
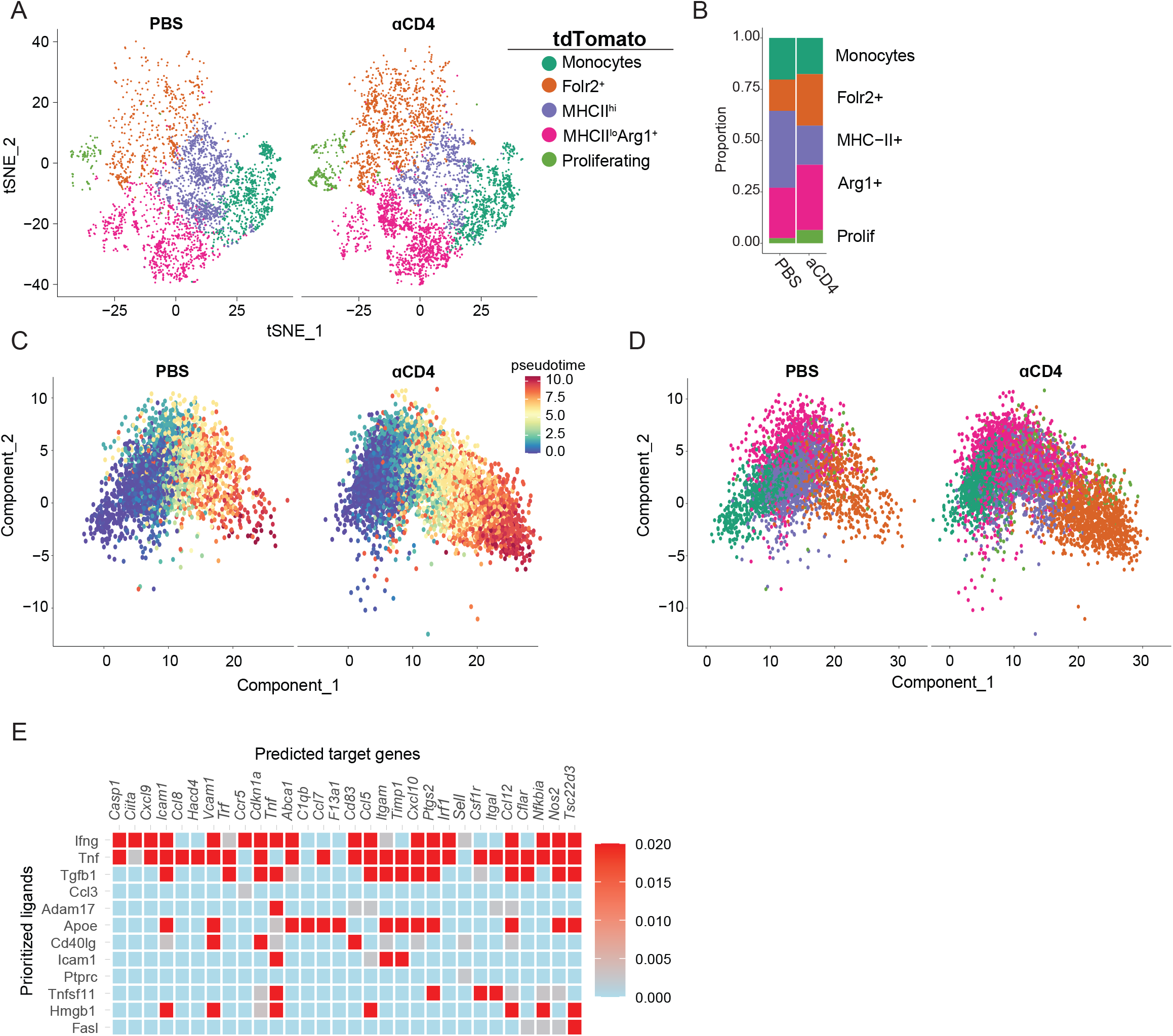
CD4 Depletion rewires monocyte differentiation toward a more protumor state. A) tSNE plot of intratumoral Tomato+ monocytes/macrophages from PBS or αCD4 treated mice on day 7. B) Proportion of each cluster among intratumoral Tomato+ monocyte/macrophages from PBS or αCD4 treated mice on day 7. C-D) Analysis of Tomato+ monocyte/macrophages using monicle3 split by pseudotime (C) and cluster (D). E) NicheNet analysis using CD4 T cells as donor cells and MHCII^hi^ macrophages as acceptor cells.

Next, we used the monocle algorithm to compare single cell differentiation trajectories of Tomato+ macrophages from PBS and anti-CD4 treated mice. Using monocytes as the root population, *Folr2*+ macrophages were predicted to be the most terminally differentiated population (Figure 4C). In the presence of CD4 T cells, cells in the monocyte cluster became primarily MHCII^hi^ or Arg1+ macrophages. However, following CD4 T cell depletion, cells from the MHCII^hi^ macrophage cluster increasingly became *Folr2*+ macrophages (Figure 4D). These data further support that MHCII^hi^ macrophages adopt a tissue resident *Folr2*+ phenotype when CD4 T cells are absent. Kinetic analysis of cluster defining genes showed that the upregulation of *Arg1* and *Folr2* was accelerated and *H2-Aa* was blunted in macrophages from CD4 depleted tumors (Supplemental figure 8). Together, the data support a phenotypic switch of monocyte fate that is dependent on the local presence of CD4 T cells.

Given our data suggests that CD4 T cells instruct monocyte differentiation into MHCII^hi^ macrophages, we utilized NicheNet^36^ to predict ligand interactions driving this fate. By setting our CD4 T cell cluster as the donor and MHCII^hi^ macrophages as the acceptor cluster, we identified several predicted ligand-receptor interactions, including *Ifng, Tnf, Tgfb1* and *Cd40l* that may drive the differentiation of MHCII^hi^ macrophages (Figure 4E). As both IFNγ and CD40 are relevant pathways that promote antitumor immunity^47^, the analysis suggest that CD4 T cells, potentially through the production of IFNγ and/or upregulation of CD40L after encountering tumor antigen, deliver these decisive signals that promote antitumor macrophage fate while mitigating monocytes from adopting a tissue-resident fate.

### Interferon gamma signaling drives antitumor TAM differentiation

Based on our NicheNet analysis, we aimed to test if MHCII^hi^ antitumor macrophage differentiation is driven by CD4-derived IFNγ, which signals through Stat1. We first assessed *Stat1* expression in TAM subpopulations from tumor-bearing wild type (WT) mice. MHCII^hi^ macrophages exhibited increased *Stat1* compared to MHCII^lo^ macrophages (Figure 5A), consistent with downstream IFNγR signaling^48^. Next, we implanted tumors into the pancreas of either WT of *Ifngr1*^*-/-*^ mice and assessed TAM phenotype at day +7 and +14. Consistent with prior work^47^, *Ifngr1*^*-/-*^ mice had significantly larger tumors at day 14 (Figure 5B) that correlated with expanded MHCII^lo^ and decreased MHCII^hi^ TAMs at both timepoints (Figure 5C). As predicted from NicheNet, tumors from *Ifngr1*^*-/-*^ mice displayed an expansion of Arg1+ TAMs (Figure 5D) and MHCII^lo^CD206^+^FRβ^+^ TAMs compared to WT (Figure 5E), suggesting that IFNγR signaling mediates TAM fate decision toward an antitumor state. However, contradictory to other models^49,50^, we found that TAM PD-L1 expression was unchanged at day 7 and only slightly reduced at day 14 in *Ifngr1*^*-/-*^ mice (Figure 5F), indicating IFNγR-independent mechanisms for driving PD-L1 expression by macrophages.

**Figure 5.**
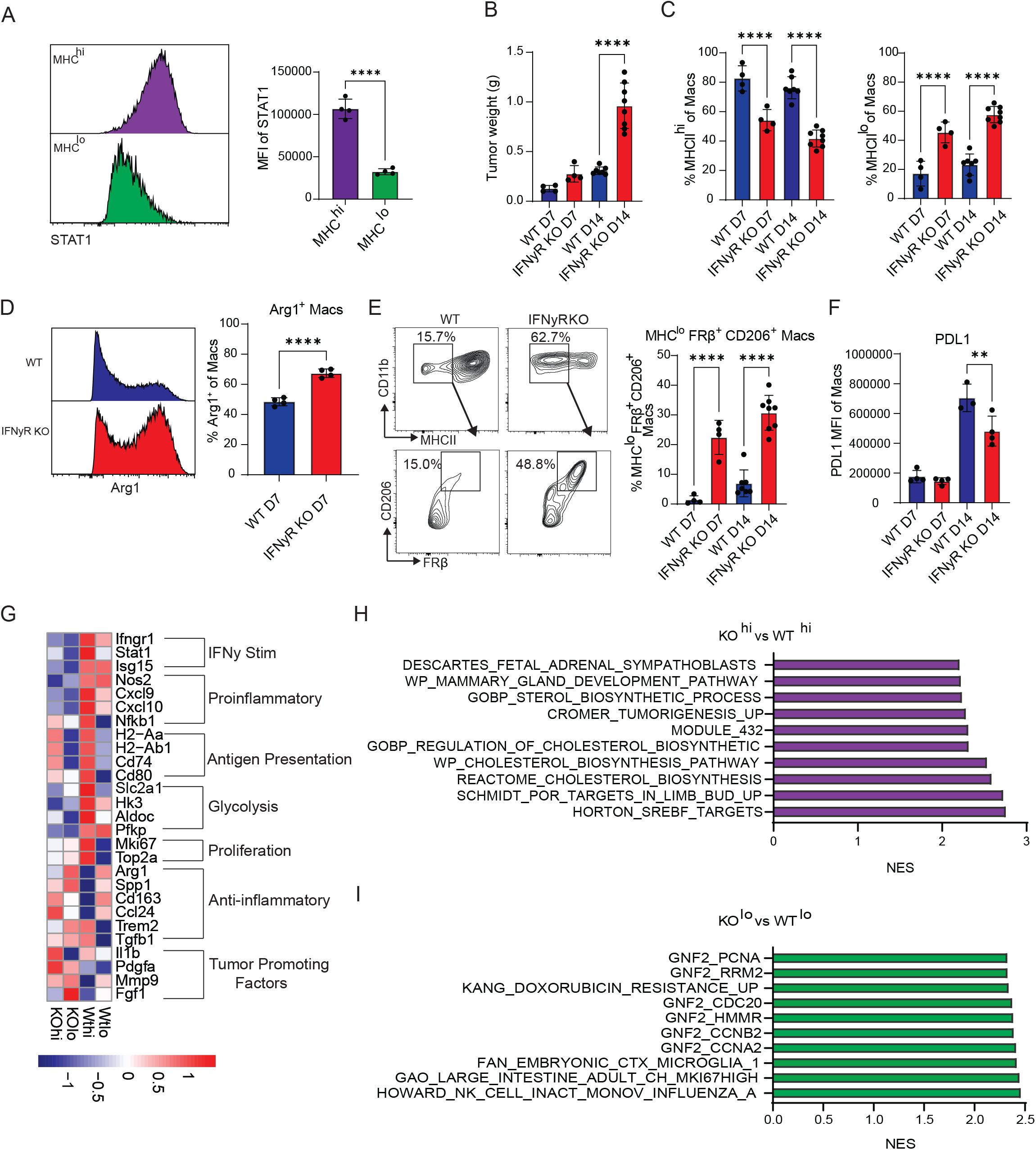
IFNγ signaling drives antitumor TAM differentiation. A) Representative Stat1 staining and Stat1 MFI in MHCII^hi^ and MHCII^lo^ intratumoral macrophages at day 7 following orthotopic *KPC2a* implantation into B6 mice. Data are mean ± S.E.M. n=4 mice per group. \*\*\**\*p*<0.0001, Student’s t-test. B) *KPC2a* tumor weights from WT and IFNyR KO mice at day 7 or 14 following orthotopic tumor implantation. n=4-8 mice per group. \*\*\*\**p*<0.0001, Student’s t-test for each time point. C) Proportion of MHCII^hi^ or MHCII^lo^ macrophages from mice in B. Each dot is an independent mouse. Data are mean ± S.E.M. n=4-8 mice per group. \*\*\**\*p*<0.0001, Student’s t-test for each time point. D) Representative Arg1 staining gated on total intratumoral macrophages and proportion of macrophages expressing Arg1 on day 7. Each dot is an independent mouse. Data are mean ± S.E.M. \*\*\*\**p*<0.0001, Student’s t-test. E) Gating strategy for detecting MHCII^lo^ macrophages and proportion of MHCII^lo^CD206+FRβ+ macrophages from mice in B. Each dot is an independent mouse. Data are mean ± S.E.M. n=4-8 mice per group. \*\*\**\*p*<0.0001, Student’s t-test for each time point. F) PD-L1 MFI gated on total intratumoral macrophages from mice in B. Each dot is an independent mouse. Data are mean ± S.E.M. \*\**p*<0.005, Student’s t-test. G) Normalized expression of selected genes following bulk RNA sequencing from sorted live CD45+CD11b+F4/80+ MHCII^hi^ or MHCII^lo^ tumor macrophages from WT and IFNyR KO mice at day 14. H-I) GSEA pathway analysis of top 300 differentially expressed genes from sorted cells in G. IFNyR KO MHCII^hi^ vs WT MHCII^hi^ mice (H) or IFNyR KO MHCII^lo^ vs WT MHCII^lo^ (I) were compared.

To elucidate transcriptomic changes driven by IFNγR signaling on TAMs, we performed RNA sequencing on sorted MHCII^hi^ and MHCII^lo^ tumor macrophages from either WT or *Ifngr1*^*-/-*^ mice (Supplemental figure 9). Differential gene expression analysis showed a downregulation in IFNγ stimulated and proinflammatory gene expression in both MHCII^hi^ and MHCII^lo^ populations in *Ifngr1*^*-/-*^ macrophage, with a compensatory increase in anti-inflammatory gene expression (Figure 5G). Moreover, *Ifngr1*^*-/-*^ macrophages exhibited a general increase in pro-tumorigenic factors such as *IL1b, Pdgfa*, and *Fgf1* which are known to play roles in metastasis and tumor growth^51–53^. We noted population dependent changes in proliferation related genes, with the MHCII^hi^ population downregulating cell cycle genes in the absence of IFNγR signaling (Figure 5G) suggesting that in contrast to well-established anti-proliferative roles for this cytokine, IFNγ can promote MHCII^hi^ macrophage proliferation in tumors. Furthermore, there was a decrease in the expression of glycolytic genes in *Ifngr1*^*-/-*^ macrophages, particularly in the MHCII^hi^ population, suggesting that IFNγ promotes MHCII^hi^ macrophage glycolysis. Pathway analysis using top differentially expressed genes comparing KO vs WT indicated upregulation of cholesterol metabolism in *Ifngr1*^*-/-*^ MHCII^hi^ macrophages, consistent with metabolic reprogramming (Figure 5H). Pathway analysis of the MHCII^lo^ subsets revealed an enrichment of processes involved in proliferation in *Ifngr1*^*-/-*^ macrophages (Figure 5I). Overall, the data suggest that IFNγR signaling is a key regulator of monocyte fate specification and has differential effects on macrophage proliferation and metabolism.

Given the proposed role of TNFα in driving inflammatory macrophages in other models^54^ and NicheNet (Figure 4E), we next assessed if loss of TNF receptor signaling impacted macrophage fate by implanting *KPC*2a tumors into TNFR1 KO mice. Although we observed a minor trend for enrichment of MHCII^hi^ macrophages, this was not significant, and the proportion and number of TAM subpopulations was largely similar at day 7 (Supplemental figure 10). Together, these data suggest that TNFR1 signaling plays a minimal role in TAM phenotype in PDA, at within least in the confines of this timepoint and model system.

### CD40/CD40L signaling drives antitumor TAM differentiation independent of IFNy

CD40 agonist has been shown to promote antitumor macrophages in the autochthonous *KPC* mouse model^13^. Signaling downstream of CD40 was enriched in fate-mapped, monocyte-derived MHCII^hi^ macrophages (Figure 4E). To determine if endogenous CD40/CD40L between CD4 T cells and macrophages promotes antitumor TAMs, we blocked this pathway using either anti-CD40L or CD40 KO mice. We found that mice lacking CD40 or anti-CD40L treatment in B6 mice had increased tumor weights compared to WT untreated controls at 7 days (Figure 6A), indicating that CD40 signaling also plays an essential role in antitumor immunity in the *KPC*2a model. Abrogating CD40/CD40L resulted in TAM populations mirroring that of CD4 T cell depleted animals, including a decrease in MHCII^hi^ macrophages, an increase of MHCII^lo^ macrophages (Figure 6B), and an expansion of both Arg1+ (Figure 6C) and MHCII^lo^ CD206+ FRβ+ (Figure 6D) macrophages. Since CD40/CD40L abrogation did not impact the frequency of CD4 T cells infiltrating tumors, the change in myeloid composition and phenotype was not merely due to an overt loss of intratumoral CD4 T cells (Figure 6E).

**Figure 6.**
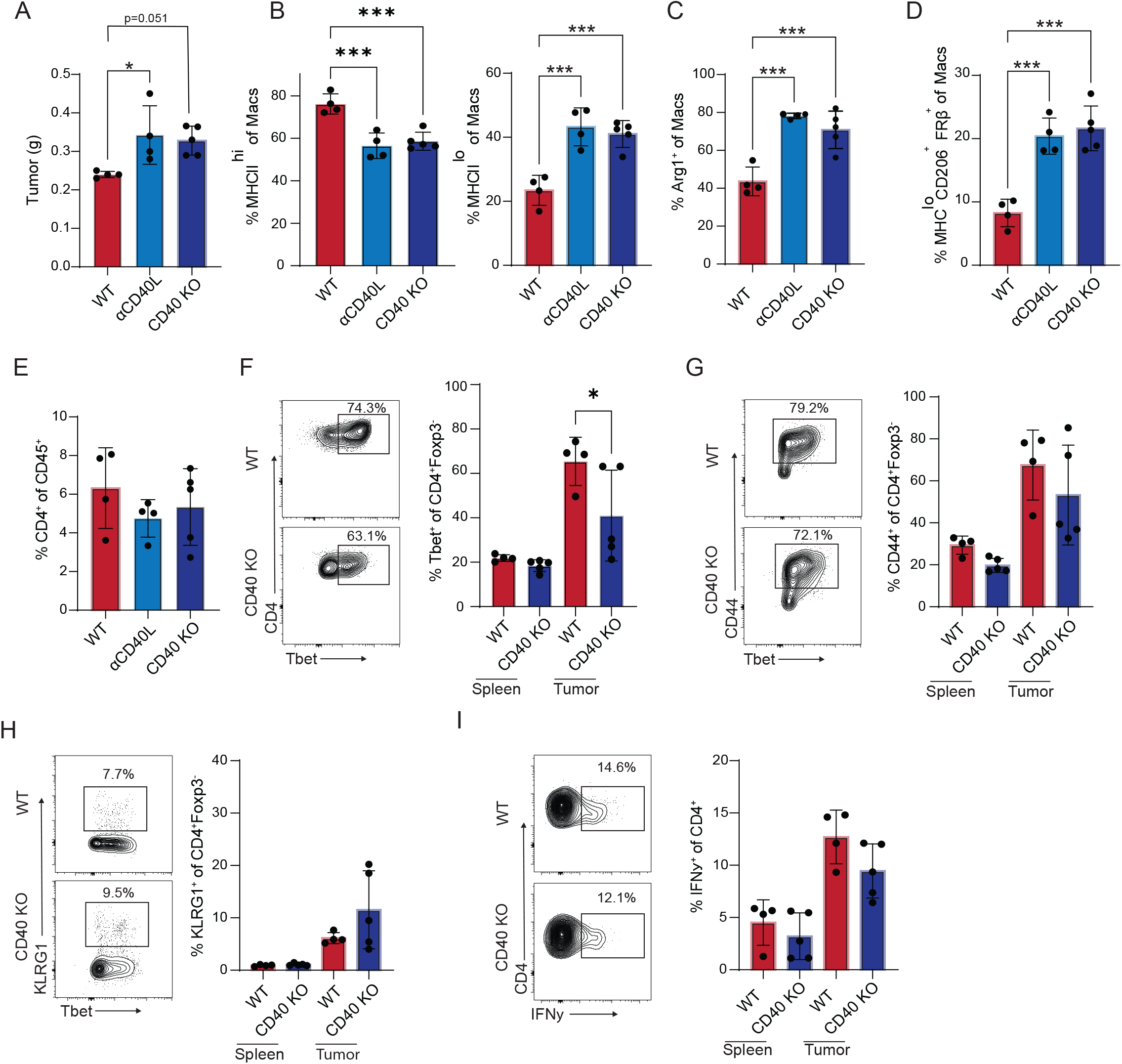
CD40/CD40L signaling drives antitumor TAM differentiation independent of IFNy. A) Orthotopic *KPC2a* tumor weights at day 7 from WT control mice, WT mice treated with CD40L blockade (αCD40L), or CD40 KO mice. B) Proportion of intratumoral MHCII^hi^ or MHCII^lo^ macrophages. Each dot is an independent mouse. Data are mean ± S.E.M n=4-5 mice per group.\*\*\**p*<0.005. One-way ANOVA with a Tukey’s posttest. C) Proportion of intratumoral Arg1+ macrophages. Each dot is an independent mouse. Data are mean ± S.E.M. n=4-5 mice per group. \*\*\**p*<0.005. One-way ANOVA with a Tukey’s posttest. D) Proportion of MHCII^lo^CD206+FRβ+ macrophages. Each dot is an independent mouse. Data are mean ± S.E.M. n=4-5 mice per group. \*\*\**p*<0.005. One-way ANOVA with a Tukey’s posttest. E) Proportion of CD4+ T cells among intratumoral CD45+ immune cells from mice in A. Each dot is an independent mouse. Data are mean ± S.E.M. n=4-5 mice per group. F) Representative plots and proportion of CD4+Foxp3-T cells that express T-bet from mice in A. Each dot is an independent mouse. Data are mean ± S.E.M. n=4-5 mice per group. \**p*<0.05, Student’s t-test for each tissue. G) Representative plots and proportion of CD4+Foxp3-T cells that express CD44 from mice in A. Each dot is an independent mouse. Data are mean ± S.E.M. n=4-5 mice per group. H) Representative plots and proportion of CD4+Foxp3-T cells that express Klrg1 from mice in A. Each dot is an independent mouse. Data are mean ± S.E.M. n=4-5 mice per group. I) Representative plots and proportion of CD4+ T cells are producing IFNy following PMA/Ionomycin. Each dot is an independent mouse. Data are mean ± S.E.M. n=4-5 mice per group.

In chronic infection models, CD40 inhibition can impair the generation of IFNγ secreting Th1 CD4 T cells^55,56^. To assess if macrophage alterations in the absence of CD40 signaling were due to impaired Th1 priming, we first assessed the activation state of intratumoral CD4 T cells in CD40 KO and WT mice. While we did see a minor reduction in the percent of T-bet+ CD4 Th1 cells in tumors from CD40 KO mice (Figure 6F), the percentage of antigen experienced CD44^hi^ CD4 T cells was similar (Figure 6G). Furthermore, loss of CD40 did not diminish the proportion of CD4 T cells that expressed Klrg1 (Figure 6H), a well-defined marker of IFNγ secreting Th1 cells^57,58^. Finally, *ex vivo* restimulation of intratumoral CD4 T cells from CD40 KO mice demonstrated a similar percentage of IFNγ producing cells compared to WT tumors (Figure 6I). Together, these data suggest that CD40/CD40L and IFNγ signaling are non-redundant pathways leading to the differentiation of monocytes into MHCII^hi^ antitumor TAMs.

### Human PDA TAM profiling

To compare our findings to human PDA, we next performed a deep characterization of myeloid cells from publicly available human tumor scRNAseq data^59^. After integrating and Seurat analysis of 6 merged human tumors, 14 distinct cell populations were identified including T-cells, B-cells, fibroblasts, monocytes and macrophages (Figure 7A, Supplemental figure 11A). We noted abundant populations of both CD4 and CD8 T cells in human PDA, similar to our scRNAseq analysis of murine orthotopic *KPC*2a tumors (Figure 3A). Clustering of just myeloid cells generated 7 unique clusters (Figure 7B). Cluster 4 expressed genes associated with undifferentiated monocytes (*S100A8, S100A9, IL1B*) while clusters 0, 1, 2, 3, 5, and 7 expressed genes of differentiated myeloid cells (*CD68, CSFR1*) (Figure 7C and Supplemental figure 11B). Cluster 0 expressed similar genes to the murine PDA Arg1+ population (*SPP1, FN1*) (Figure 7C), but lacked *ARG1* expression. Cluster 3 expressed genes associated with antigen presentation (*HLA-DQB2*), paralleling the murine intratumoral MHCII^hi^ macrophages. Cluster 3 also expressed *CD1C* and *CD1D* (Figure 7C), consistent with a phenotypical relationship to monocyte derived dendritic cells (MoDCs) which have been well-defined in humans^60^. Notably, *CD1C+HLA-DQB2+* Cluster 3 was enriched for genes such as *CD14* and *CSF1R* which are well-established markers on cells of monocyte lineage^60^ (Supplemental figure 11B). Cluster 1 expressed high levels of *FOLR2* (Figure 7C,D) resembling tissue resident macrophages in the *KPC*2a mouse from our studies above, and in human breast cancer^45^. Both Clusters 6 and 2 expressed higher levels of *TREM2* (Figure 7D), which has been proposed to be a marker of monocyte derived TAMs^45^. Cluster 2 expressed higher transcript levels of complement associated genes like *C1QB*, while Cluster 6 expressed higher levels of *GPNMB* (Figure 7C, D), supporting heterogeneity among *TREM2+* TAMs. Next, we performed pseudotime analysis to interpret single cell differentiation trajectories of monocytes-to-macrophages in human PDA. Using monocytes as the origin point, this model predicted two endpoint trajectories, either to *FOLR2*+ and *TREM2*+ TAMs or MHCII^high^ MoDCs terminal lineages (Figure 7E).

**Figure 7.**
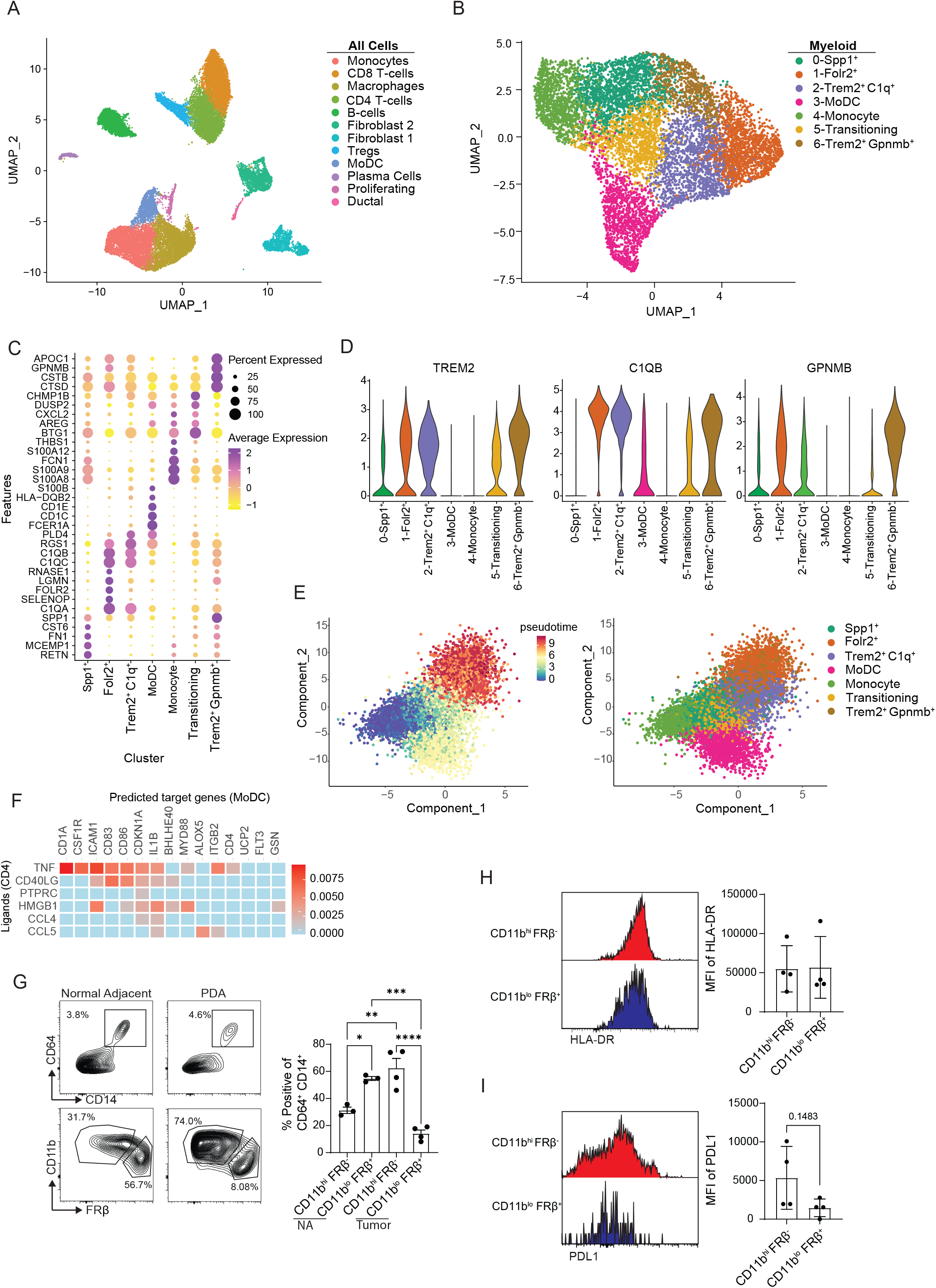
Profiling of TAMs from Human PDAC. A) UMAP clustering of 6 merged human tumors from Elyada *et al*. Cell types were identified based on top differentially expressed genes as described in Materials and Methods B) UMAP re-clustering of myeloid clusters in A. MoDC, monocyte-derived dendritic cell. C) Heatmap of normalized gene expression of the top differentially myeloid genes from clusters in B. D) Violin plots of selected genes that drive monocyte/macrophages subclusters in B. E) Pseudotime of myeloid clusters depicted in (B) using mocicle3 split by pseudotime and by cluster F) NicheNet analysis to CD4 T cells as donor cluster and MoDCs as acceptor cluster. G) Gating scheme and frequency of myeloid subpopulations from resected human PDA (n=4) and normal adjacent (n=3) tissue. Gated on Live, CD15^-^ cells. Data are mean ± S.E.M. \**p*<0.05, \*\**p*<0.005, \*\*\**p*<0.001, \*\*\*\**p*<0.0001. One-way ANOVA with a Tukey’s posttest. H) Representative HLA-DR staining and HLA-DR MFI from CD11b^hi^ FRβ- and CD11b^lo^ FRβ+ macrophages (CD64+ CD14+) from human PDA and normal adjacent tissue. Gated on live, CD15^-^ cells. Data are mean ± S.E.M. I) Representative PDL1 and PDL1 MFI by CD11b^hi^ FRβ- and CD11b^lo^ FRβ+ macrophages (CD64+ CD14+) isolated from human PDA and normal adjacent tissue.

Given that our mouse data suggested that CD4 T-cells instruct monocyte differentiation toward a more immunostimulatory phenotype, we next predicted potential interactions between CD4 T cells and MoDCs using NicheNet. This algorithm predicted that TNF and CD40/CD40L signaling were enriched between these two populations (Figure 7F). Unlike mouse data, IFNγR signaling was not predicted as a regulator between these two populations, potentially due to the markedly low levels of *IFNG* transcripts in advanced human PDA (Supplemental figure 12).

Finally, to examine the relative contributions of different TAM subsets to human PDA, we performed flow cytometry on samples from resected PDA and normal adjacent human pancreas. We identified two major macrophage populations within both the normal adjacent tissue and tumor: CD11b^hi^ FRβ^-^ and CD11b^lo^ FRβ^+^(Figure 7G). Given that embryonically derived macrophages express lower levels of CD11b^61^, we suspect that CD11b^lo^ FRβ^+^ are bonified tissue resident TAMs. Consistent with this, most macrophages in normal adjacent tissue adopted a CD11b^lo^ FRβ^+^ phenotype, however within the tumor more macrophages were CD11b^hi^ FRβ^-^ (Figure 7G), Thus, these data are consistent with the majority of the TAM pool in human PDA being derived from monocytes. Unlike in our mouse model, FRβ^+^ TAMs expressed similar amounts of HLA-DR compared to FRβ^-^ cells (Figure 7H), consistent with Folr2 expressing TAMs in human breast cancer^45^. However, like the *KPC*2a orthotopic PDA model, FRβ^-^ macrophages expressed PDL1 to a higher extent (Figure 7I). These data suggest that monocyte derived CD11b^hi^ FRβ^-^ may play more of a role in regulating immune responses and may reflect a lack of a sustained functional CD4 Th1 cell population in the chronic human disease. Together, these data support lineage identity between monocyte- and tissue resident-derived TAM populations in human PDA.

## Discussion

The goal of immunotherapy is to generate antitumor T cells capable of infiltrating into the TME and eliciting potent antitumor function through the production of cytotoxic molecules, expression of costimulatory proteins such as CD40L, and effector cytokines such as IFNγ. While understanding how the TME shapes antitumor T cells has been a major focus, we investigate here how antitumor T cells shapes the TME. We focus on myeloid cells because they are abundant in solid tumors, have plasticity, and can be complicit in tumor pathogenesis. We identify that tumor specific CD4 T cells, but not CD8 T cells, shape monocyte differentiation and promote antitumor macrophage fate decisions through nonredundant IFNγR and CD40 pathways. Notably, in the absence of a productive tumor specific CD4 T cell response, a subset of monocyte-derived macrophages adopt a phenotypic and transcriptomic profile similar to that of tissue resident macrophages supporting that monocytes can masquerade as tissue-resident TAMs in a poorly immunogenic tumors. Thus, just as mechanisms of ‘*adaptive resistance’* such as PD-L1 are at play^62^, when a productive T cell response is engaged, mechanisms of ‘*adaptive assistance’* contribute to immune-mediated control of solid tumors. We posit that further investigation into adaptive assistance mechanisms, in which mononuclear phagocytes are a key target, will inform the development of effective immune-based strategies for cancer patient treatment.

We identify that the TAM pool is mostly derived from circulating monocytes in neoantigen+ PDA. A landmark macrophage fate mapping study in neoantigen negative PDA models showed that that a subset of embryonically-derived pancreatic resident macrophages are maintained throughout adulthood and expand during tumorigenesis^23^. The embryonically-derived tissue resident macrophages were suggested to account for half of the macrophage pool and display distinct phenotypes/functions from monocyte derived TAMs^23^. In contrast to previous fate mapping studies in PDA, we identified that most TAMs are from monocyte derived precursors rather than tissue resident pancreas cells. Recent studies support that the presence of tumor specific T-cells can accelerate monocyte recruitment^28^, which could potentially outcompete resident macrophages and account for differences in relative contribution. Thus, this discrepancy may be due to the antigenicity of the tumor model, and a source of further investigation. Using a specific monocyte fate mapping approach, we validate the premise that both monocyte and embryonically derived TAMs are present in PDA and have distinct phenotypes, even in the presence of tumor specific CD4 T cells. However, our data also suggest that in the absence of the CD4 T cell-induced microenvironmental changes, monocytes adopt a tissue resident phenotype. It is important to note that even in the absence of a neoantigen, orthotopic *KPC* tumors still have significant CD4 T cell infiltration^63^, which could account for distinct phenotypes of tissue resident and monocyte derived TAMs.

Our data support that local environmental cues impacting monocyte fate rather than stochastic cell-intrinsic fate choices underlies macrophage heterogeneity. While CD8 T cells are spatially located near TAMs and engage in an antigen specific manner that promote T cell exhaustion^28^, we show that CD4 T cells are a key supplier of inflammatory cues within the TME that promote monocytes differentiation into pro-inflammatory macrophages. Critically, pro-inflammatory macrophages express antigen presentation machinery, secrete proinflammatory cytokines and promote antitumor responses and appear broadly similar to macrophages primed in the autoimmune milieu of the pancreas of the non-obese diabetic mouse^64^. CD4-TAM interplay has been seen in other models where infusion of tumor reactive T cells lead to upregulation of inflammatory markers on TAMs^65^, however it was unclear if this was a repolarization of macrophages or changes in tumor infiltrating monocyte fate decisions. Furthermore, Th2 CD4 T cells can skew macrophage phenotypes toward a more anti-inflammatory state within tumors^66^, highlighting the linkage between macrophages and CD4 T cells. Licensing of macrophages by CD4 T cells does not seem to be restricted to the TME as effector CD4 T cells can exacerbate autoimmune pathology by engaging myeloid cells through TNFR signaling^67^. Given our findings with CD40L blockade and CD40-deficient mice, CD4 T cells likely engage newly recruited monocytes physically, and that these physical interactions in context of the TME cytokine milieu dictate monocyte fate specification. Overall, our study reveals that tumor specific CD4 T cells shape the TME and supports those therapeutic efforts such as engineered T cell therapies should focus on stimulating Th1 responses for enhancing clinical benefit.

Our fate mapping approach suggests that monocytes can differentiate into antitumor (MHCII^hi^) or protumor (MHCII^lo^) macrophages in the TME. To the best of our knowledge, no previous study has performed in-depth mapping of monocyte differentiation in cancer. Our data using CCR2 reporter mice supports the premise that monocyte differentiation within inflamed tissue is heterogeneous and based off local environmental cues rather than a sequential predetermined pathway. Similar to our model, studies utilizing human PDA have found that monocytes differentiate into either immunostimulatory monocyte-derived DCs or immunosuppressive TAMs^68^. Therefore, the identification of factors that can interfere with the differentiation of monocytes into the MHCII^lo^ macrophage or promote the differentiation of monocytes into the MHCII^hi^ subset represent key therapeutic targets. In addition, tissue-resident macrophages appear predisposed to adopt a protumor phenotype and contribute to tumor growth in PDA^23,69^, as well as other cancer models^70,71^. These observations suggest that tissue-resident macrophages may be particularly resistant toward rewiring to an immunostimulatory state which warrants further investigation. A recent report describing macrophage polarization as type 1 diabetes progresses in autoimmune-prone NOD mice may provide phenotypic clues in this regard^64^.

Previous reports from our lab have found that IFNγR signaling on host cells is required for responses to immunotherapy in PDA^47^. Furthermore, CD40 agonist has shown to be transiently effective as a monotherapy in the *KPC*2a tumor model of PDA^14,31^ and has shown promising results clinically^72^. However, what cell types and through what pathways these mediators act on to promote antitumor immunity is still largely unclear. Through this work, we show that IFNyR and CD40 signaling rewires monocyte differentiation toward an antitumor state which correlates with improved outcomes. These findings are directly relevant clinically and suggest that targeting both pathways could improve antigen presenting functions of TAMs and boost T cell-mediated antitumor immunity. Overall, we mapped monocyte differentiation in PDA and found that adaptation of pro-tumor or antitumor TAM state is driven by tumor-specific CD4 T cells and that in the absence of these cells, monocytes can adopt a tissue-resident state. These findings have implications as to the design of immunotherapies and support that inclusion of CD4 T cells during adoptive cell therapy will be critical for antitumor activity in solid tumors.

## Acknowledgements

We acknowledge the University of Minnesota Flow Cytometry Resource for technical assistance and the University of Minnesota Research Animal Resource (RAR) staff for animal husbandry and veterinary services. M.T.P. is supported by an American Heart Association Predoctoral fellowship #903380. A.L.B is supported by a computational training award from the American Association of Immunologists. MF was supported by the Ministry of Science and Higher Education of the Russian Federation (Agreement No. 075-15-2022-301). KZ was supported by Priority 2030 Federal Academic Leadership Program. J.W.W. is supported by NIH R01 AI165553-01A1 and American Heart Association CDA #855022. I.M.S. is supported by NIH R01 CA249393, R01 CA255039, and P01CA254849, Department of Defense #PA200286, an American Association for Cancer Research (AACR), Pancreatic Cancer Action Network Career Development Award (17-20-25-STRO), an AACR Pancreatic Cancer Action Network Catalyst Award (19-35-STRO), American Cancer Society Institutional Research Grant (124166-IRG-58-001-55-IRG65), Randy Shaver Cancer and Community Fund and has pilot awards from the Masonic Cancer Center and Cancer Research Training Initiative (University of Minnesota Medical School).

## Author Contributions

M.T.P., I.M.S. and J.W.W designed the research. M.T.P. and A.L.B. contributed to the study design. M.T.P., A.L.B, Y.X., Z.C.S., P.R.S., A.E.K., E.A.M., E.C.H. performed experiments. M.T.P., A.L.B., and Y.X. analyzed data, interpreted data, and prepared figures and tables. Y.X., M.M.F and K.Z. performed analyses of the single-cell RNA sequencing data. M.T.P. and A.L.B wrote the manuscript, and all authors revised the manuscript.

