## Supplementary Figures 1-12 for "Tumor-specific CD4 T cells instruct monocyte differentiation in pancreatic ductal adenocarcinoma"

**Patterson et. al., Supplementary Figures 1-12**

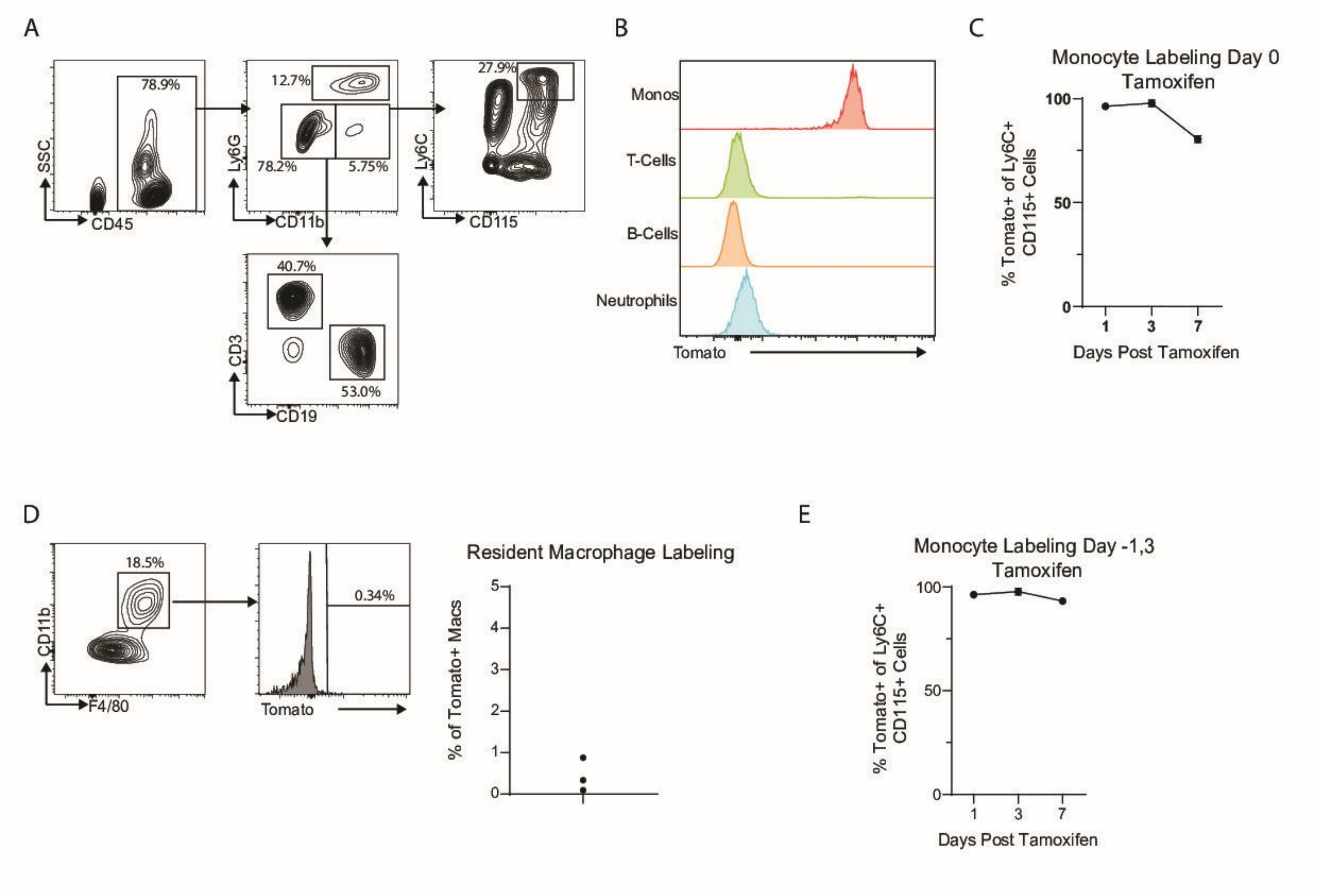


**Supplementary Figure 1. Labeling efficiency of CCR2$^{CreER}$ R26$^{TdTomato}$ monocyte fate mapping mice.**
A) Gating strategy for analysis of circulating blood immune cells from tumor bearing CCR2$^{CreER}$ R26$^{TdTomato}$ mice 1 day post tamoxifen administration.

B) Tomato reporter expression by circulating blood immune subsets from tumor bearing CCR2$^{CreER}$ R26$^{TdTomato}$ mice 1 day post tamoxifen administration.

C) Proportion of circulating blood monocytes that are Tomato+ from tumor bearing CCR2$^{CreER}$ R26$^{TdTomato}$ mice treated with tamoxifen on day of tumor implantation (Day 0) (n=4). The data are mean +/- S.E.M.

D) Labeling and quantification of pancreatic resident (Tomato-) macrophages from non-tumor bearing CCR2$^{CreER}$ R26$^{TdTomato}$ 1 day post tamoxifen administration. CD11b+F480+ macrophages were gated on live, CD45+Ly6G- cells.

E) Proportion of circulating blood monocytes that are Tomato+ from tumor bearing CCR2$^{CreER}$ R26$^{TdTomato}$ treated with tamoxifen 1 day prior to tumor implantation (Day -1) and on day 3 after tumor implantation (n=4).

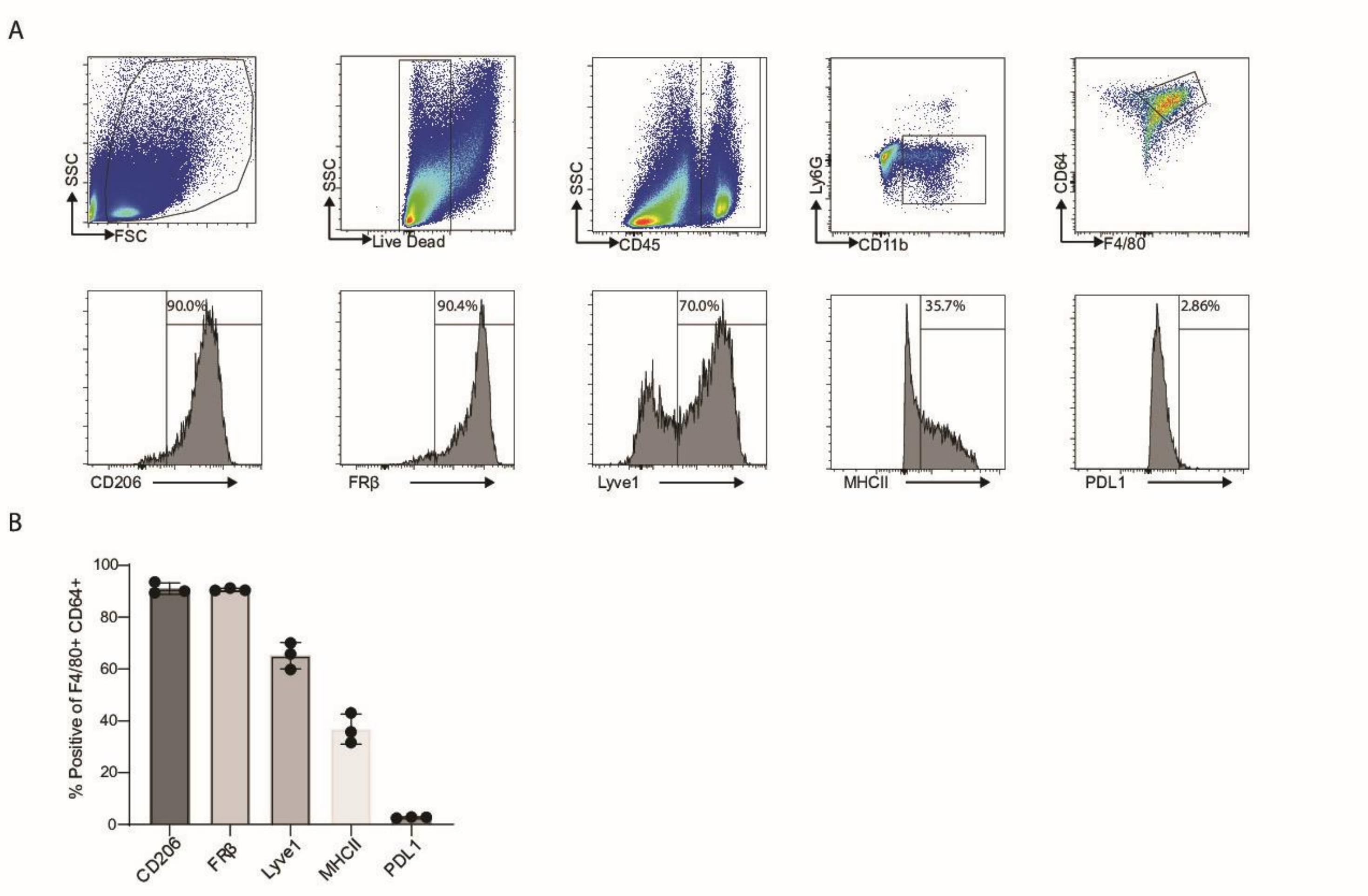


**Supplementary Figure 2. Phenotyping of steady state pancreatic resident macrophages.**
A) Gating strategy for analysis of pancreatic macrophages from untreated and non-tumor bearing $CCR2^{CreER}$ $R26^{TdTomato}$ mice. Live cells are gated off single cells.

B) Proportion of CD64+F4/80+ pancreatic macrophages that express the indicated marker (n=3). Each dot is an independent mouse. Data are mean ± S.E.M.

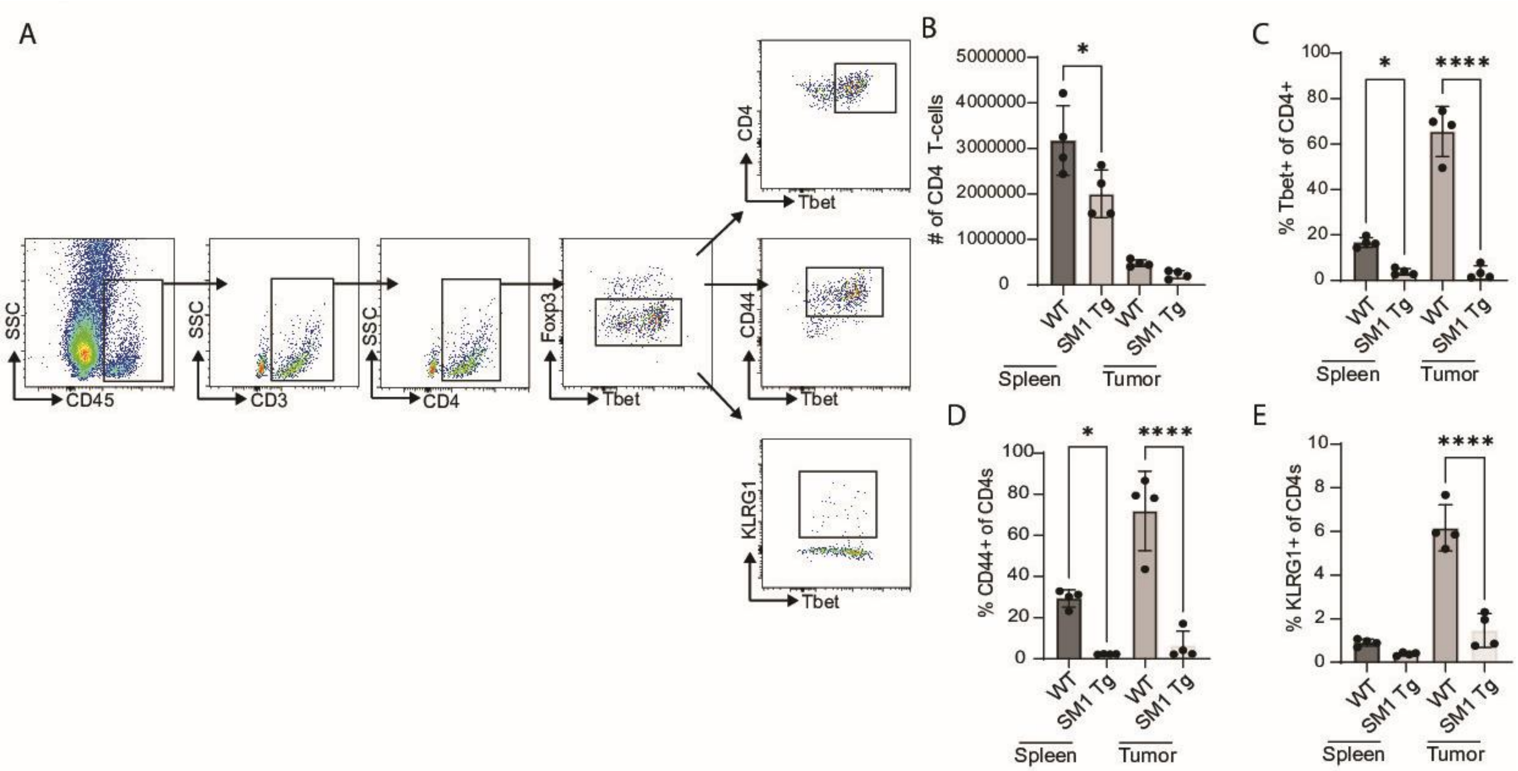


**Supplementary Figure 3. CD4 phenotyping in absence of Neoantigen specific T-cells**
A) Gating strategy for analysis of CD4 T cells from WT mouse on day 7 post orthotopic tumor implantation. Representative plots are gated off live, single cells.

B) CD4+T cell counts from WT or SM1xRAGKO mice at day 7 post tumor implantation. (n=4 per group). Each dot is an independent mouse. Data are mean ± S.E.M. *$p<0.05$, Student's t-test for each tissue.

C-E) Proportion of CD4+Foxp3- (Tcons) T cells that express Tbet (C) CD44 (D) or KLRG1 (E) in WT or SM1xRAGKO mice at day 7 post tumor implantation. (n=4 per group). Each dot is an independent mouse. Data are mean ± S.E.M. *$p<0.05$, **$p<0.005$, ****$p<0.0001$, Student's t-test for each tissue.

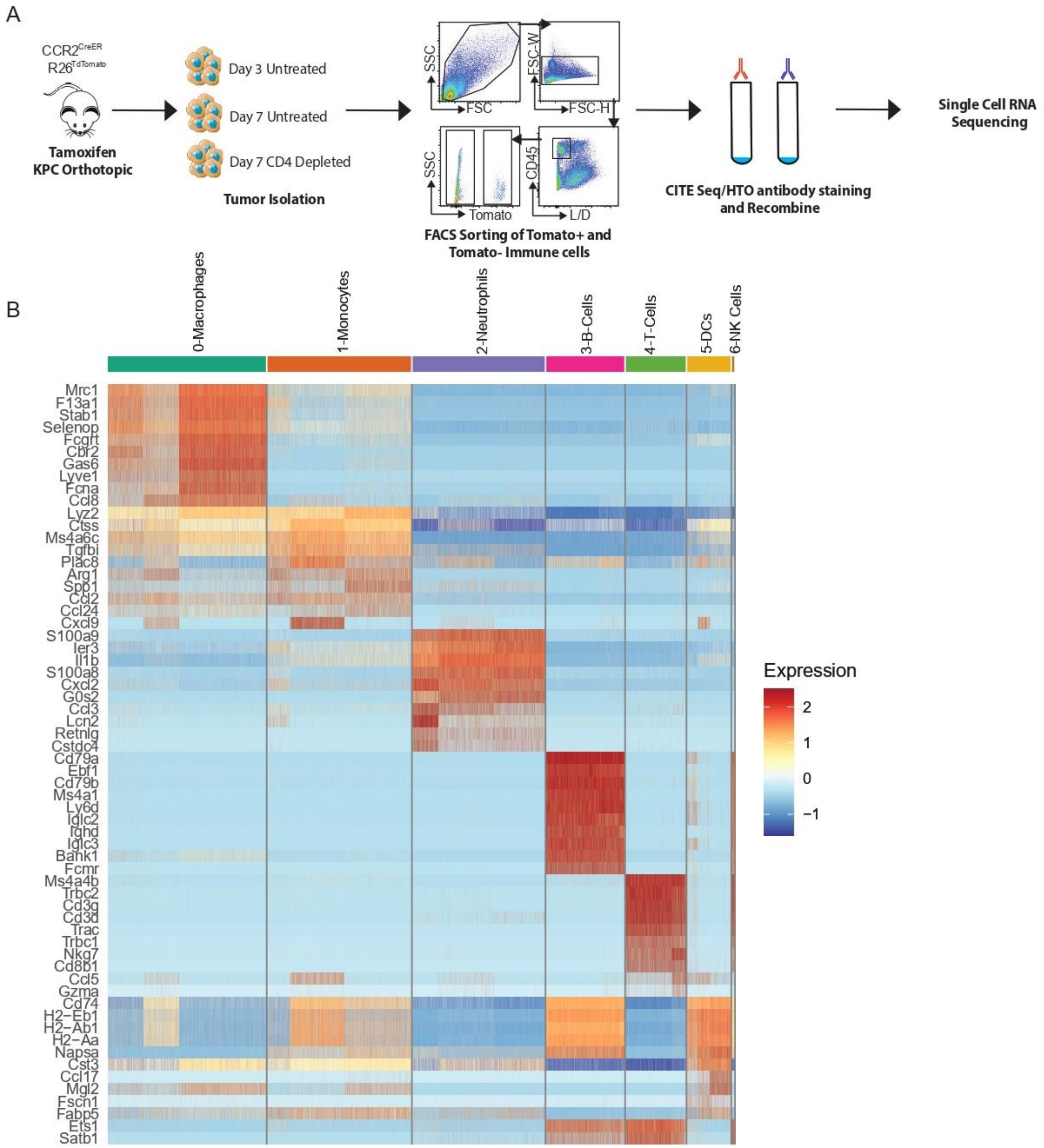


**Supplementary Figure 4. scRNAseq monocyte fate mapping in murine PDA**

A) Experimental approach for scRNAseq analysis of immune cells infiltrating *KPC*2a tumors. CCR2$^{CreER}$ R26$^{TdTomato}$ mice were orthotopically implanted with *KPC*2a tumors. One cohort was treated with anti-CD4 at day -1 and day +2. All three cohorts were administered tamoxifen on the day of tumor implantation. Tumors were isolated from a total of 4 mice per cohort per timepoint (day 3 and day 7). Tumors from mice treated with anti-CD4 were harvested on day 7. Tumor single cell suspensions were generated and Tomato+ and Tomato- immune cells were FACS sorted, labeled with CITE-Seq antibodies and hash tagged and pooled at a 1:1 mixture for scRNAseq.

B) Heatmap showing top 5 differentially expressed genes for each cluster. singleR was used to name cell populations based on top differentially expressed genes.

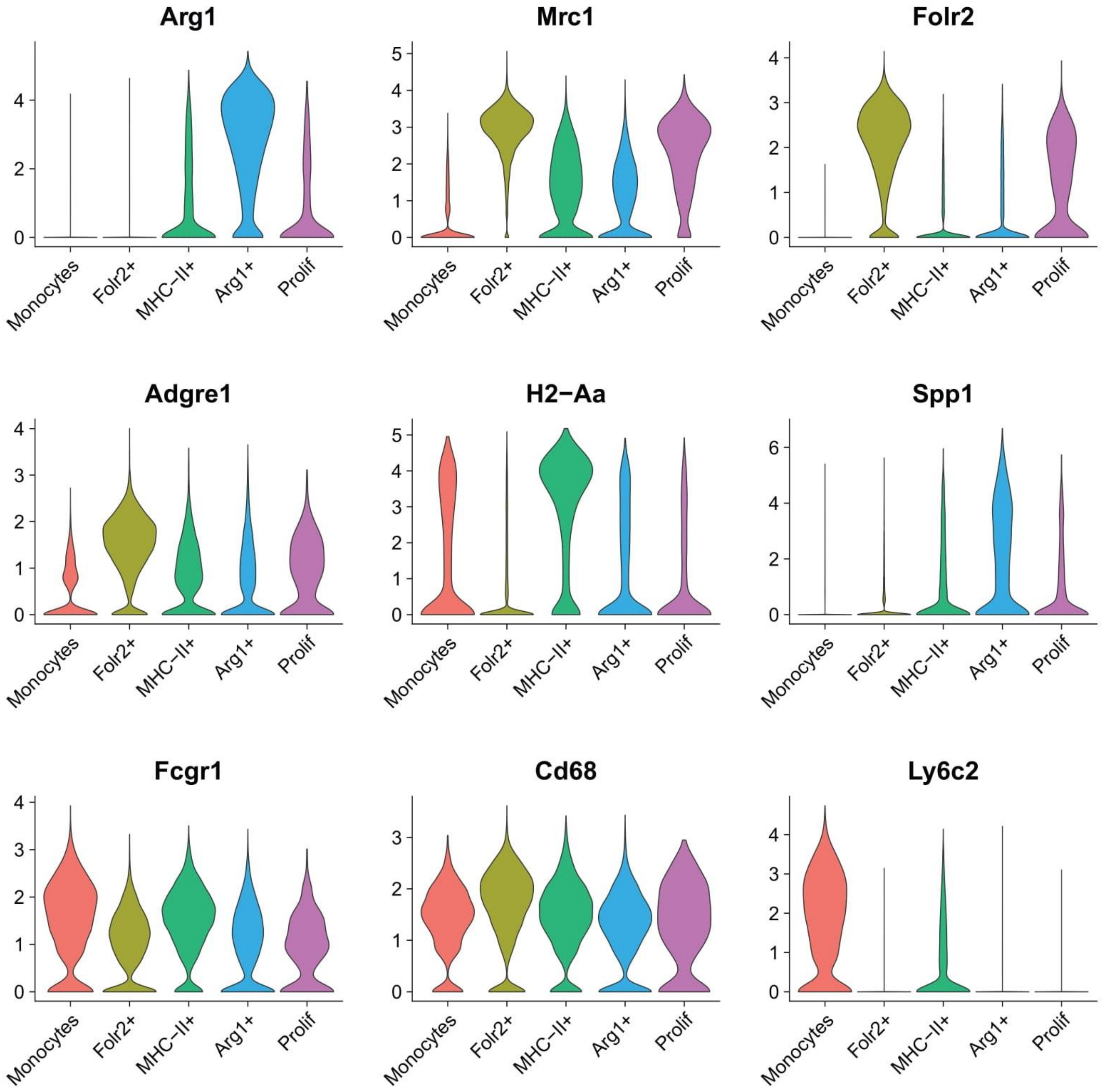


**Supplementary Figure 5. Monocyte/macrophage cluster defining genes.** Violin plots of selected cluster defining genes from scRNAseq data.

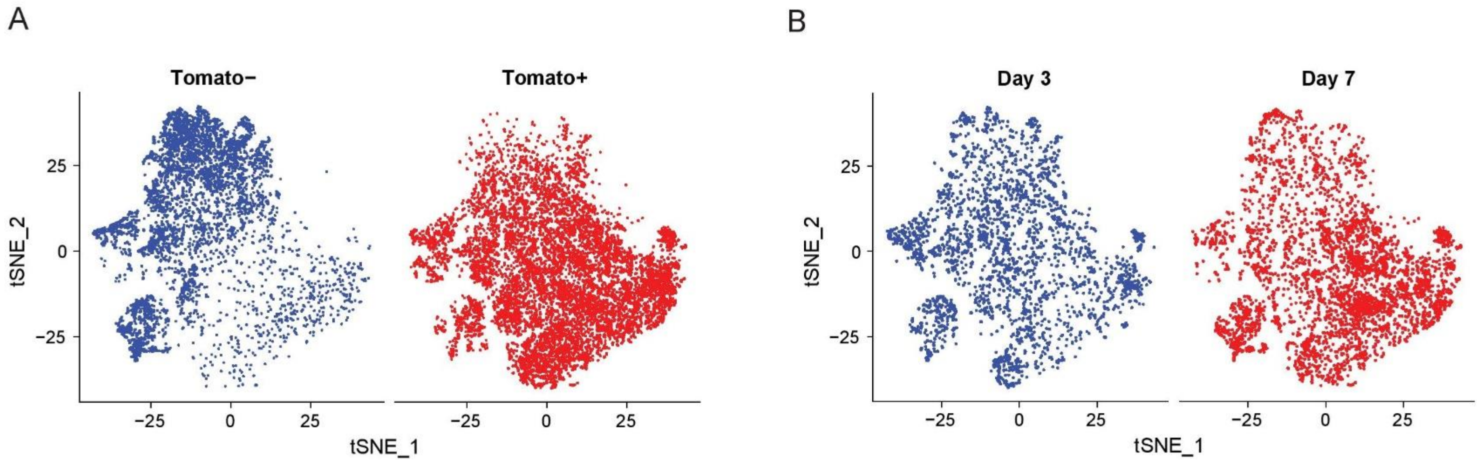


**Supplementary Figure 6. scRNAseq fate mapping temporal changes**
A) tSNE plot showing Tomato+ and Tomato- cells from Day 3 and Day 7 untreated and proportion of each monocyte/macrophage cluster among Tomato+ and Tomato- myeloid cells.
B) tSNE plot and proportion of each monocyte/macrophage cluster at day 3 and day 7 untreated.

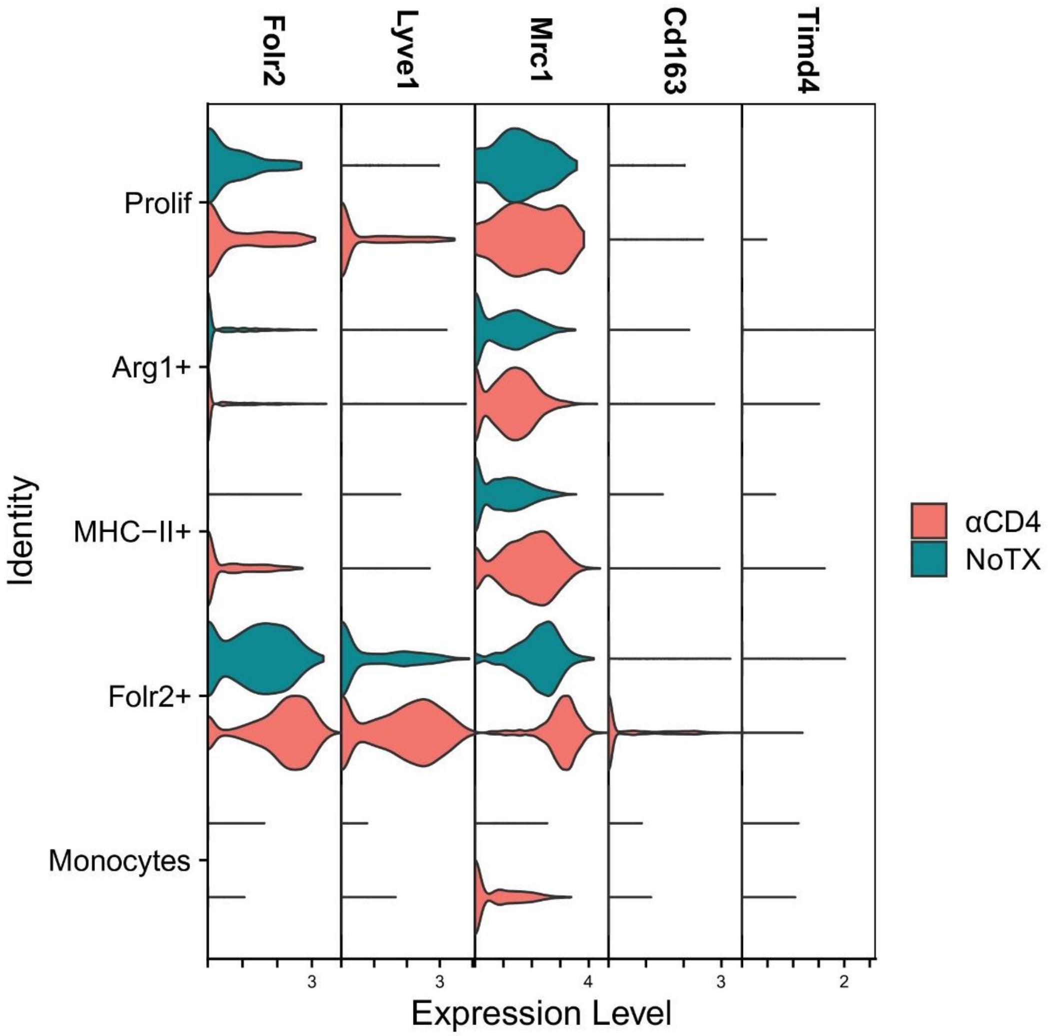


**Supplementary Figure 7. Expression of tissue resident genes in monocyte derived macrophages.** Violin plots of selected genes associated with a tissue resident phenotype. Genes are stratified by Tomato+ monocyte/macrophage cluster from untreated (NoTX) or anti-CD4 treated (αCD4) *KPC*2a tumor bearing mice on day 7 posttumor.

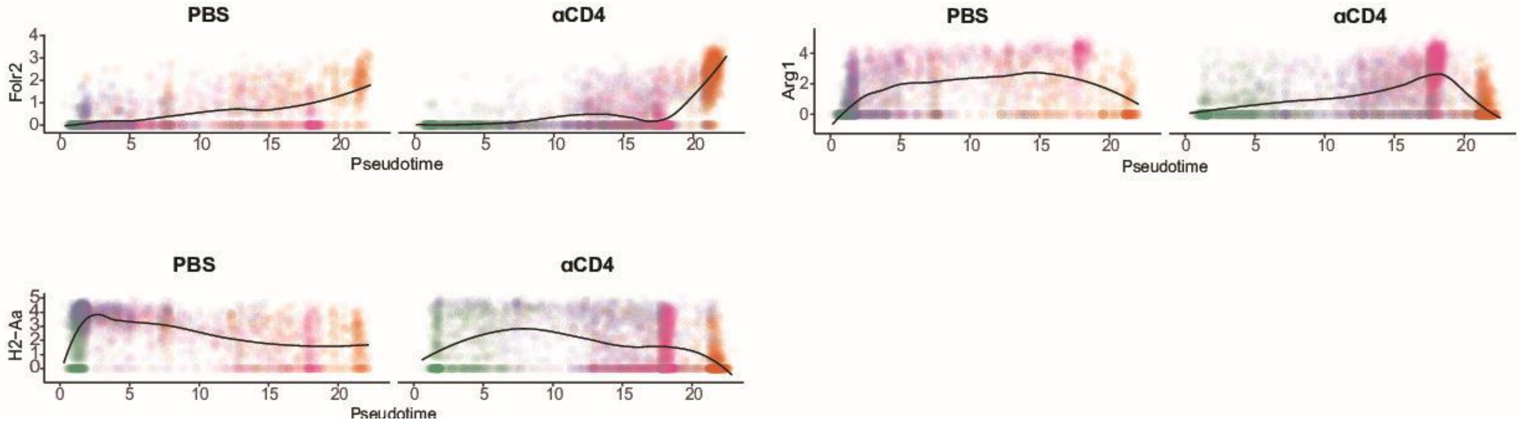


**Supplementary Figure 8. Kinetic analysis of cluster defining genes.** Kinetic analysis of cluster defining gene expression over pseudotime. Folr2 defines Folr2+ cluster, MHCII defines MHCII+ cluster and Arg1 defines Arg1+ cluster.

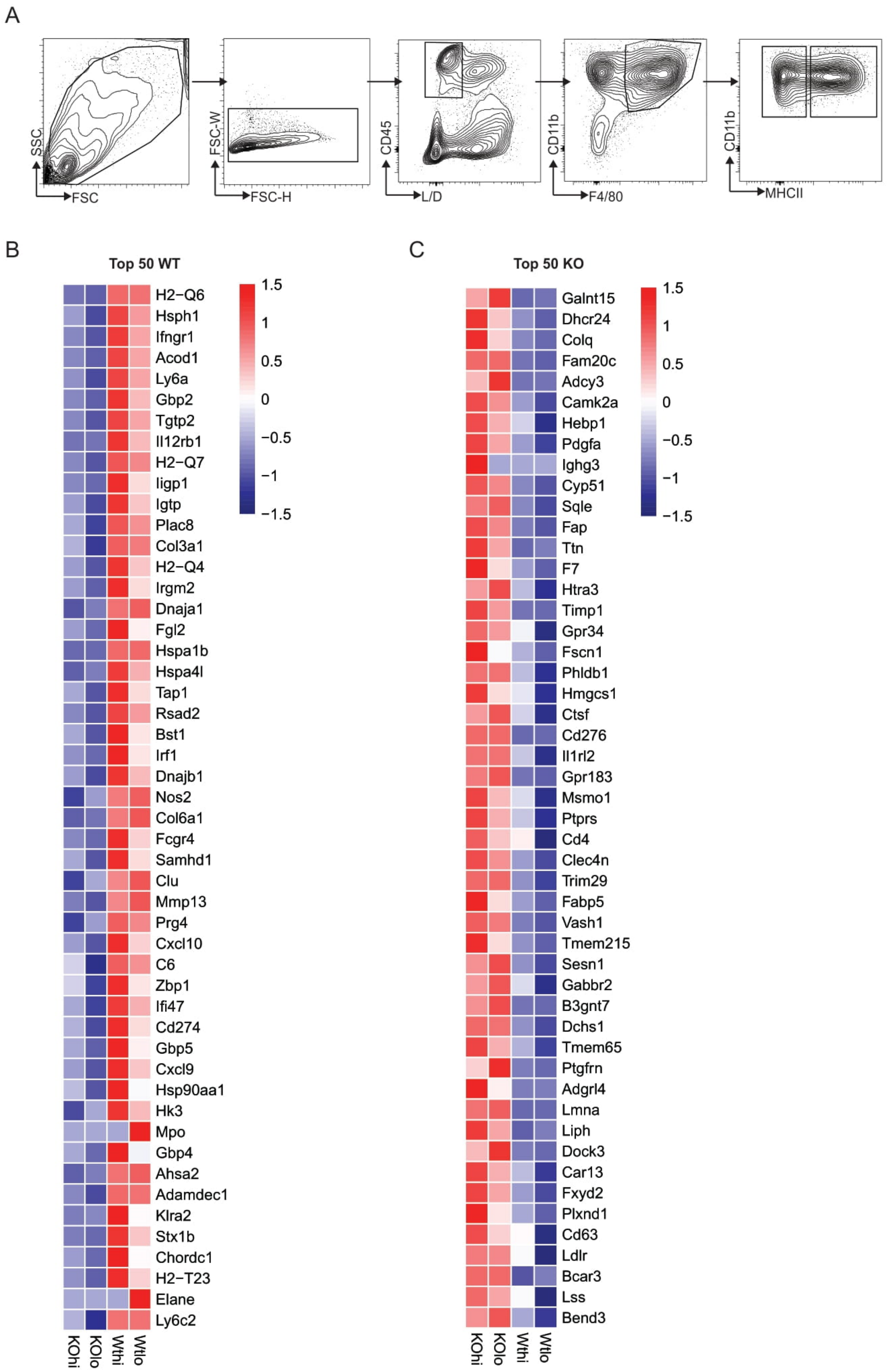


**Supplementary Figure 9. Bulk RNA sequencing of TAMs from IFNyR KO and WT mice.**
A) FACS sorting strategy for isolation of MHCII$^{hi}$ and MHCII$^{lo}$ TAMS from 4 pooled WT and 4 pooled IFNyR KO mice on 14 days post tumor implantation.

B) Heat map of top 50 differentially expressed genes for each macrophage population.

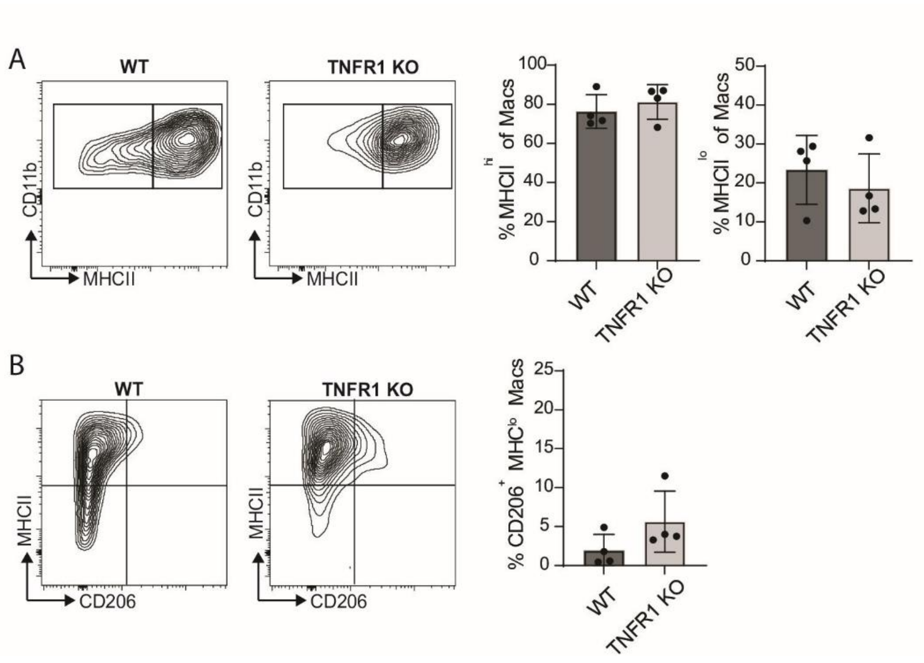


**Supplementary Figure 10. Impact of host cell Tnfr1 deletion on macrophage phenotype.**
A) $MHCII^{hi}$ vs $MHCII^{lo}$ macrophage frequency from tumors isolated from WT and TNFR1 KO mice 7 days after implantation (n=4 mice per group). Populations are gated on CD64+ F4/80+ cells. Each dot is an independent mouse. Data are mean ± S.E.M.

B) $MHCII^{lo}$ CD206+ macrophage frequency in tumors from WT and TNFR1 KO mice in A. Populations are gated on CD64+ F4/80+ cells. Each dot is an independent mouse. Data are mean ± S.E.M.

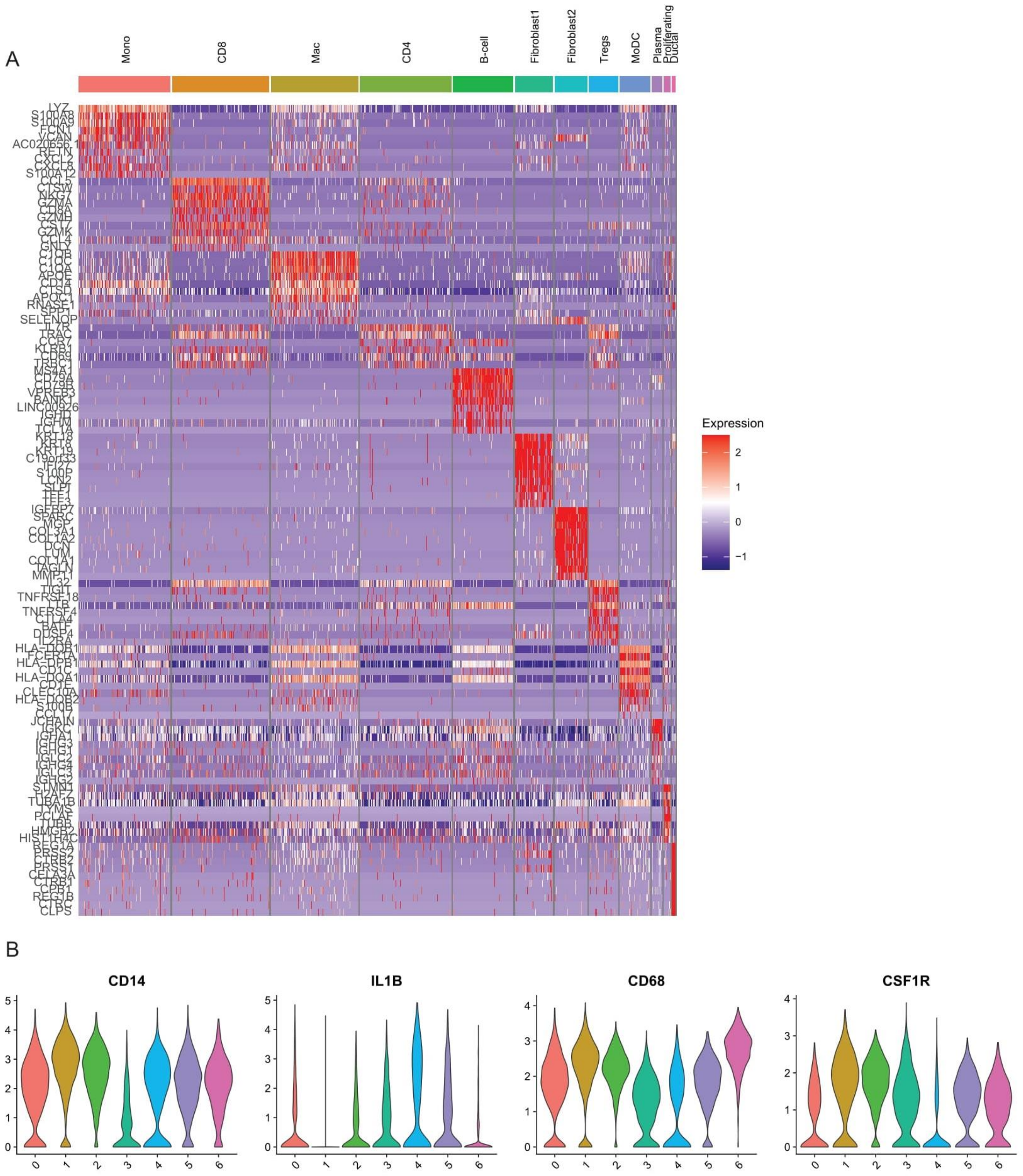


**Supplementary Figure 11. Human PDA scRNAseq analysis.**
A) Heatmap of top differentially expressed genes from 6 resected human PDA tumors from Elyada *et al*. Cell populations were clustered in the UMAP space and named based off their top differentially expressed genes

B) Violin plots of selected cluster defining genes from A.

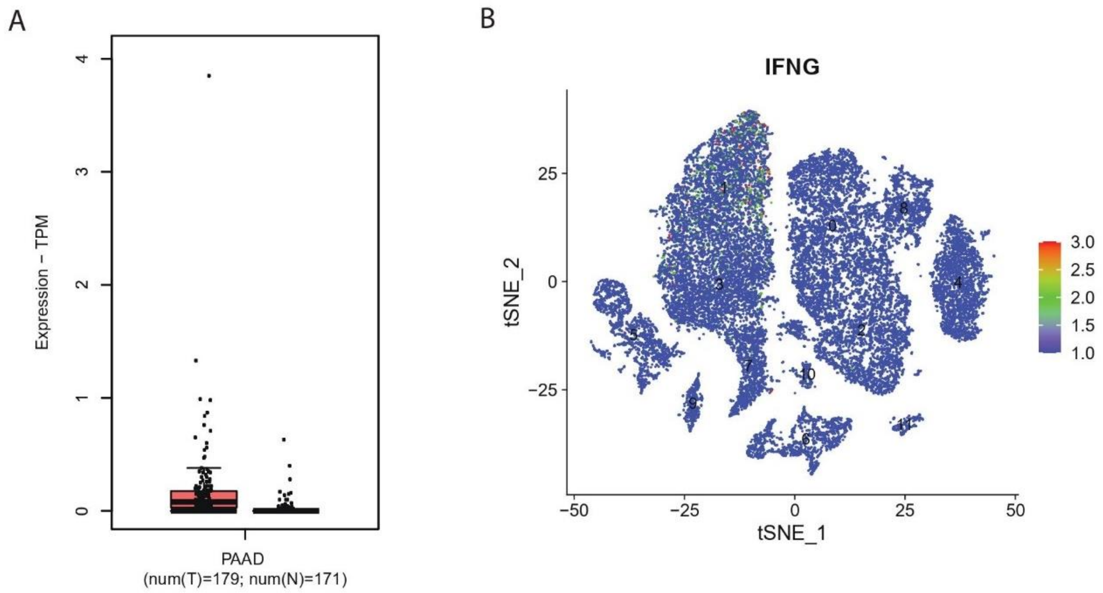


**Supplemental Figure 12.** Lack of *IFNG* in resected human PDA.
A) Log fold change expression of *IFNG* from human PDA (e.g., PAAD) from tumors (red) and normal adjacent (grey) was analyzed using GEPIA.

B) tSNE plot of *IFNG* overlaid on scRNAseq data from 6 merged human PDAs from Elyada *et al*.
